# Emergence and influence of sequence bias in evolutionarily malleable, mammalian tandem arrays

**DOI:** 10.1101/2022.07.13.499775

**Authors:** Margarita V Brovkina, Margaret A. Chapman, Matthew L. Holding, E. Josephine Clowney

## Abstract

The radiation of mammals at the extinction of the dinosaurs produced a plethora of new forms—as diverse as bats, dolphins, and elephants—in only 10-20 million years. Behind the scenes, adaptation to new niches is accompanied by extensive innovation in large families of genes that allow animals to contact the environment, including chemosensors, xenobiotic enzymes, and immune and barrier proteins. Genes in these “outward-looking” families are allelically diverse among humans and exhibit tissue-specific and sometimes stochastic expression. Here, we show that outward-looking genes are clustered in tandem arrays, enriched in AT-biased isochores, and lack CpG islands in their promoters. Models of mammalian genome evolution have not incorporated the sharply different functions and transcriptional patterns of genes in AT-versus GC-biased regions. To examine the relationship between gene family expansion, sequence content, and functional diversification, we use population genetic data and comparative analysis. First, we find that AT bias can emerge with gene family expansion *in cis*. Second, human genes in AT-biased isochores or with GC-poor promoters experience relatively low rates of *de novo* point mutation today but are enriched for functional variants. Finally, we find that isochores containing gene clusters exhibit low rates of recombination. We hypothesize that the depletion of GC bases in outward-facing gene clusters results from tolerance of sequence variation and low recombination. In turn, high AT content exerts a profound effect on their chromatin organization and transcriptional regulation.

## Introduction

Reports of newly sequenced genomes frequently describe gene families that have “bloomed,” undergoing explosive diversification in the focal species (1–3). Gene blooms are expansions *in cis* that result in arrays of dozens or even hundreds of genes. During gametogenesis, these tandem duplication events are thought to arise via incorrect crossovers between paralogues or via non-homologous repair of chromosome breaks (4). The resulting expansions can confer unique life history traits recognized as definitive characteristics of the species: Examples include Cytochrome p450 genes for plant detoxification in koala and insects, lipocalins for pheromone communication in mouse, NK cell receptors for viral response in bats, keratins for whale baleen, venom production in snakes, and amylase copy number for starch consumption in modern humans (1,2,5–11). The definitive gene family of mammals, the caseins, arose through local duplication of enamel genes (12). In the mouse, we have shown that genes in copy-number-variable blooms exhibit extremely high AT content in their promoters and are often located in AT-biased regions of the genome (13).

GC content in mammalian genomes varies markedly at the megabase scale (14). Since the earliest days of cytology, variation in staining patterns of DNA-binding dyes were apparent across the nucleus (heterochromatin and euchromatin) or along chromosomes (banding patterns). Banding patterns served as the original genetic maps and allowed scientists to link genetic phenotypes to physical positions in DNA (15). Banding patterns were found to reflect local variation in AT/GC content: Giemsa-staining “G bands” are AT-biased, and Quinacrine-staining “Q bands” are GC-biased (15–18). Early reports suggested that G-bands were depleted for genes; that genes in G-bands tended to be “tissue-specific;” and that genes in Q-bands tended to be “housekeeping genes” (15,19). Moreover, human-chimp divergence rates were found to be higher in AT-rich G-bands than in GC-rich Q bands (20). In the genome sequencing era, breaks between bands were found to correspond to local transitions in GC content, and bands were found to be composed of smaller “isochores” with locally consistent GC content (21,22). While isochore definition has been debated, a representative classification breaks the human genome up into ∼3000 isochores of 100kb-5Mb that range from 35-58% GC (22–25).

The variation in GC content along the chromosome that is observed in mammals is not a general feature of metazoan, animal, or even vertebrate genomes. Both average GC content and the amount of local variation show wide divergence across clades (26,27), leading to adaptationist speculation that isochore structure serves a function related to endothermy (28). However, consensus has emerged that GC-biased gene conversion (gBGC) following meiotic recombination, which occurs in most eukaryotes, is one important contributor to isochore emergence in mammals (26,29). In this process, crossovers are statistically more likely to result in gene conversion towards more GC-rich sequences, resulting in a higher likelihood of inheriting higher-GC alleles. As stated by Pouyet and colleagues, “The gBGC model predicts that the GC-content of a given genomic segment should reflect its average long-term recombination rate over tens of million years” (30,31). In this model, the isochores themselves do not serve an adaptive function, but rather have emerged due to the molecular genetic (“neutral”) forces of meiosis. Over evolutionary time, the GC-increasing effect of recombination is counteracted by the AT-increasing effect of point mutation due to mutation of fragile cytosines to thymines (32–37). As the rate of recombination and cytosine loss may themselves be influenced by sequence context (i.e. recombination more likely in GC-rich regions, cytosine mutation more common in AT-rich regions), positive feedback could have caused large genomic regions to diverge (20,30,32,38). However, the influence of these neutral forces on the genes contained within AT-versus GC-biased isochores has not been described.

Here, we examine the local and isochore-level AT/GC content of human protein-coding genes. Genes located in AT-rich regions of the genome have unique and consistent characteristics: They are copy-number-variable families located in tandem arrays, are expressed in terminally differentiated cells, are cell surface or secreted proteins, lack CpG islands in their promoters, and often have stochastic or variegated expression. These protein families are overwhelmingly involved in the “input-output” functions of an organism: sensation of the environment, protection from the environment, consumption of the environment, and production of bodily fluids. AT-versus GC-skewed isochores differ in their replication timing and histone marks, they associate in nuclear space with other isochores of the same type, and they occupy different domains within the nucleus (16,39–41). The distinct treatment of AT-versus GC-rich isochores by the molecular machinery of the mammalian cell means that the genes located in AT-rich isochores must experience distinct molecular events from those located in GC-rich isochores.

Next, we ask how mammalian genes with outward-looking functions came to be located in AT-rich regions of the genome. By comparing gene blooms of different sizes within the human genome and across mammalian species, we find that AT content is not necessarily inherited from the ancestral species but can emerge with cluster expansion. Many of the mutagenic and repair processes that contribute to the neutral mutation spectrum are likely to differ in strength across AT-rich and GC-rich regions (reviewed in (42)). Using human population genetic data, we analyze allelic variation, patterns of point mutation, and recombination in human genes located in AT-versus GC-biased isochores. We find that genes in paralogous clusters are subject to less recombination than genes located near non-paralogues. Recombination may be dangerous in gene clusters due to the potential for chromosome rearrangements or within-cluster ectopic exchange that leads to duplications or deletions. It could also separate genes in large families from locus control regions they depend on for expression.

In addition, we find that while genes in AT-biased isochores have high sequence diversity among humans and divergence across species, they currently exhibit relatively low rates of *de-novo* point mutation, as expected from the neutral mutation spectrum. Instead, they appear to have accumulated sequence variants over evolutionary time. These genes also lack CpG islands in their promoters. We hypothesize that the functional roles of genes whose protein products interface with unpredictable and rapidly changing molecules in the environment (“outward-looking genes”) make them particularly likely to tolerate point mutations. Lack of purifying selection on these genes is expected to shift GC content down over evolutionary time due to deamination of cytosine leading to C->T transitions (42); intolerance of recombination would prevent gBGC from shifting GC content back up (37). We propose a model in which reduced recombination and relaxation of selection on point mutations act together to strand arrays of outward-looking paralogues in wells of low GC content (Figure 1). Loss of CpG islands and residence in AT-rich genomic regions predisposes these genes to exotic forms of highly tissue-specific transcriptional regulation (43–47).

**Figure 1:**
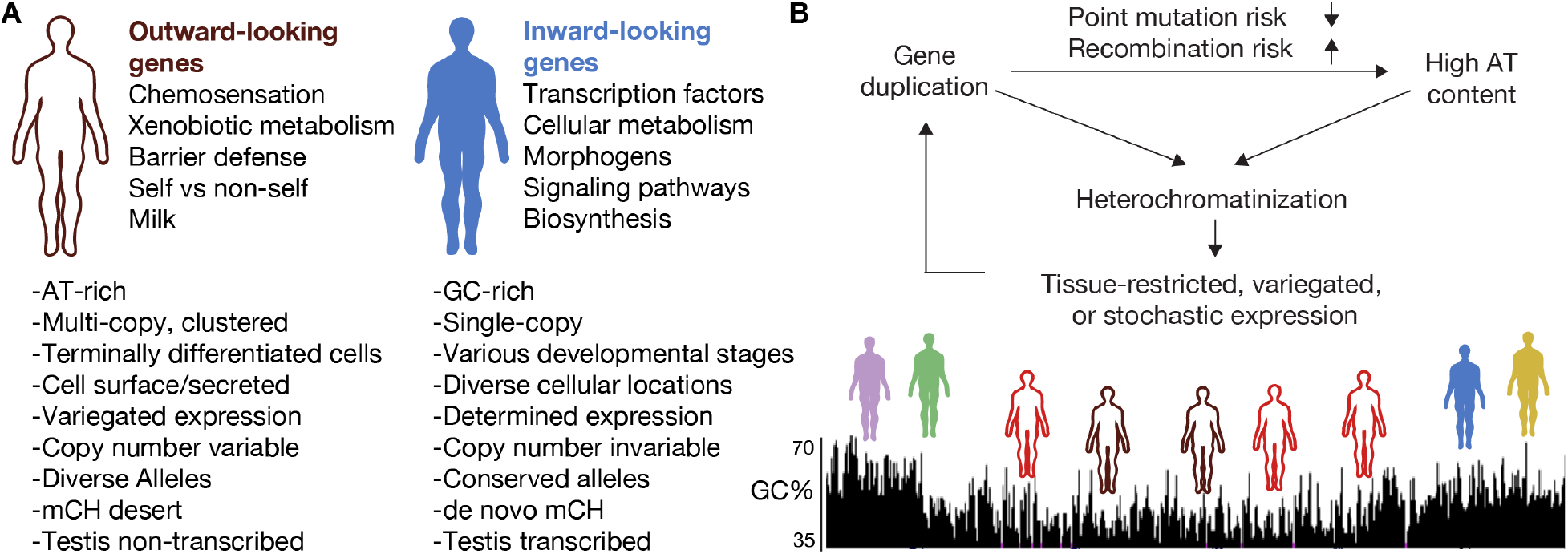
Outward- versus inward- looking genes. (A) Summary of inward-looking and outward-looking gene families and their genomic distinctions. This table is inspired by (48). Descriptions derive from our analyses here and from references cited throughout the text. (B) Model of possible relationships between expansion of gene families in cis, genomic architecture, selective forces, and mode of expression.

## Results

### Characterizing a set of human isochores and their gene contents

We first comprehensively characterized the types of genes located in AT- versus GC-biased isochores and assessed if genes in AT-biased isochores are more likely to have numerous paralogues as neighbors. Isochore boundaries have been computed for previous genome assemblies, including for hg38 (22–24,49). While previous methods of isochore annotation binned the genome into 100kb pieces or used manual inspection to annotate isochore ends, we used a segmentation algorithm which detects transitions in GC content and created a UCSC Genome Browser track for easy visualization of isochore assignments versus other genomic elements (50). Cozzi et al compare various methods of isochore assignment, including GC Profile, showing that differences are subtle and are most prevalent in the mid-ranges of GC content. We tested three resolutions and selected a map that visually matched 100kb-Mb transitions in GC content using the %GC track on the UCSC Genome Browser (Fig 2A,B). At this resolution, we call 4328 isochores; these range from ∼30-70% GC and most are between 100kb and 5Mb (Figure 1C, D, S2A-C, Supplemental Tables 1 and 2).

**Figure 2:**
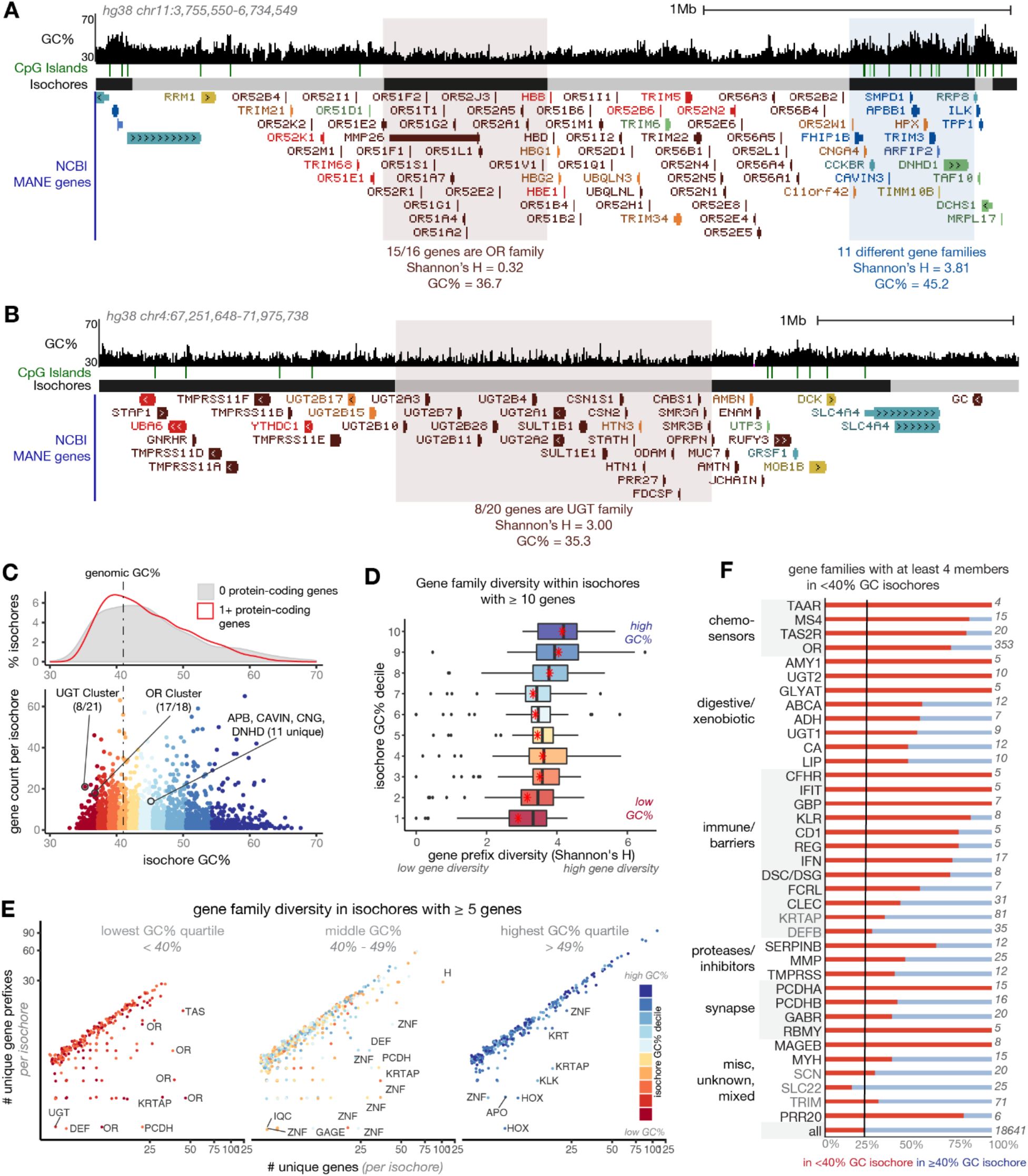
AT-rich isochores in the human genome contain tandem arrays of genes with outward-looking functions. (A, B) UCSC Genome Browser screenshots showing GC% trajectory (53), CpG islands (54), our isochore calls, and simplified gene models. Gene models are colored according to k-means clusters described below. Gene name prefix diversity (Shannon’s H) is shown for the highlighted isochores. (C) Relationship between isochore GC% and gene contents. Isochores shown in A, B are highlighted. Colors show how we binned isochores into deciles according to GC%. Dashed line shows the mean GC content of the human genome (41%). (D) Boxplots representing gene name diversity (Shannon’s H) of gene-rich isochores across GC% deciles shown in (C). Red points indicate the mean prefix diversity of the decile; black lines show medians. Here and throughout, most groups are statistically different from one another, except for adjoining groups; full statistical comparisons are presented in Supplemental Table 3. (E) Comparison of number of genes in an isochore versus number of different gene name prefixes represented in that isochore, for isochores of at least five genes.

We next defined the home isochore of each gene in the NCBI MANE set. The MANE set includes one promoter and splice isoform for each intact, protein-coding gene and omits pseudogenes and complex “gene parts” such as V, D, and J segments of the B and T cell receptors (51). On average, higher-GC isochores were more likely to contain protein-coding genes and had higher gene density (Figure 2C, Figure S2A, B) (15). To compare features across isochores, we ordered isochores by GC% and divided them into ten groups (deciles) of ∼400. To test whether tandemly arrayed “gene blooms” were associated with AT-rich isochores, we used Shannon’s H to measure gene name prefix diversity in isochores with at least ten genes (Figure 2D). Gene names were least diverse in AT-rich isochores, consistent with the presence of tandem arrays in these isochores. We extended our analysis to isochores with fewer genes (at least five) and again found that gene arrays were less common in high-GC isochores (Figure 2E, S2H,I).

While AT-rich isochores are longer, gene diversity is lower in AT-rich isochores across the length distribution and for isochores with different numbers of genes (Figure S2F, G). Arrays in AT-rich isochores contained more paralogues, and the kinds of proteins present in AT-rich arrays versus GC-rich arrays were different (Figure 2E, S2F): Arrays in AT-rich isochores often served sensory, digestive, or barrier functions, while arrays in higher-GC isochores most often contained *HOX* and *ZNF* transcription factors or arrays of histones (Figure 2E, S2I).

Only 25% of genes are located in isochores <40% GC. What sorts of gene families have bloomed in these isochores? As GO analysis is biased by the annotations that are available, especially for extremely tissue-specific genes, we simply searched for common gene name prefixes in isochores <40% GC (Figure 2F). Gene families with at least four members in high-AT isochores were overwhelmingly involved in chemosensation (*OR, TAAR, TAS2R, MS4*); xenobiotic metabolism (e.g. *AMY1, UGT2, ADH*); and immune, defense, and barrier functions (e.g. *KLR, IFN, DSC/DSG*). Human ORs were also shown previously to be located in AT-biased isochores (52). Examples of high-GC isochores with diverse gene members and high-AT isochores with repetitive gene members are shown in Figure 2A, B and Figure S2D,E. While immunoglobulin parts do not appear in the MANE gene set, arrays of immunoglobulin V regions are also highly AT-biased (Figure S2G). These analyses demonstrate that AT-biased regions of the human genome contain tandem arrays of genes with outward-looking functions. We invite readers to explore these patterns further on our TrackHub: http://genome.ucsc.edu/s/mbrovkin/hg38.

Each dot is an isochore; isochores falling below the trend line contain clusters of genes with the same prefix. Isochores are labeled by the most common gene name prefix in that isochore. (F) Gene families with at least four members in isochores <40% GC. 25% of all genes are in isochores less than 40% GC (black line). Red bars depict proportions of genes with that prefix that are located in <40% GC isochores. Gene family prefixes shown in black text are enriched in AT-rich isochores, while those shown in grey text (e.g. *KRTAP*, *TRIM*, *SCN*) have multiple family members in AT-rich isochores but are not enriched there. Functions of these gene families are marked at left, and total number of genes with that prefix in the MANE set are shown at right.

### Categorizing human genes according to local patterns of AT/GC content

We show above that tandemly arrayed genes serving outward-looking functions are enriched in AT-rich regions of the human genome, as they are in mouse (13). We sought next to ask whether the regulatory and transcribed regions of genes, which occupy a fraction of genome space, also vary in GC content across different kinds of genes, and whether GC content of local gene features follow that of the isochore context. The null expectation of this analysis is different from a statistical genetic versus a molecular point of view. From a statistical standpoint, the expectation would be that GC% would co-vary between gene parts (e.g, flanking, promoter, coding sequence) and their isochore context. From a molecular point of view, transcribed units and regulatory regions would be expected to have a GC content aligned with their regulatory or protein-coding function, and would not necessarily match their genomic location. We calculated GC% in 50bp sliding windows along the transcriptional unit (transcription start site to transcription end site, TSS-TES) and 1kb flanking regions for genes in the MANE set (51,53). Our analysis here includes introns, but clustering on exons and flanking regions produced similar results (Figure S3B). We used iterative k-means clustering to group the 18,640 MANE genes into 3x3 sets (Figure 3A, S3A). The top-level clusters (1-, 2-, 3-) reflect overall differences in AT content in different genes (Figure S3A), while the subclusters (1.1, 1.2, 1.3 etc, Figure 3A) reflect variation in AT content of the promoter, transcriptional unit, and 3’ region. To capture both the broad isochore context of genes and their local sequence features, we use both the isochore AT/GC metric and the local sequence-based k-means clustering throughout this study; each gene in the MANE set is assigned uniquely to one home isochore and one k-means cluster (isochore deciles 1-10, red-blue palette; k-means clusters 1.1-3.3, rainbow palette). Cluster and isochore assignments and other gene-linked data are provided in Supplemental Tables 4 and 5.

**Figure 3:**
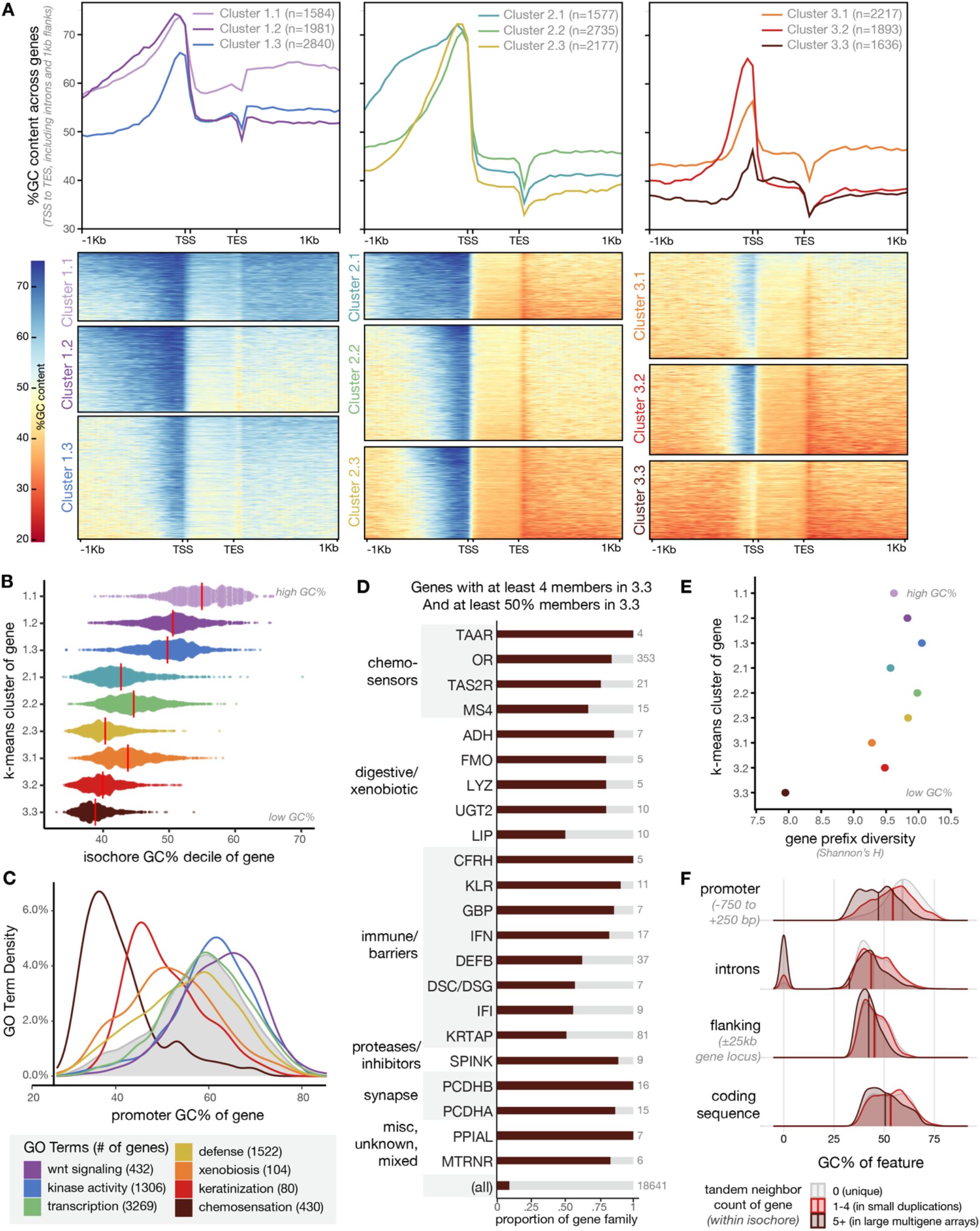
Genes with outward-looking functions have high local AT content. (A) GC content trajectory for human protein-coding genes in the MANE set. Genes were subdivided by iterative k-means clustering. At top, the average GC content trajectory for each k-means cluster is shown as a line graph. At bottom, each gene is a row and GC content across the transcriptional unit and flanking regions is depicted from red (high AT) to blue (high GC). Rainbow colors assigned to each k-means cluster here will be used throughout. (B) Relationship between k-means cluster assignment and home isochore GC% for each gene in the MANE set. Red lines depict medians. (C) GO term distribution by promoter GC content for genes in the MANE set. Genes with immune, barrier, chemosensory, and xenobiotic functions have AT-skewed promoters. Genes with developmental and intracellular functions have GC-skewed promoters. Grey shading shows promoter GC content distribution of the whole MANE set. (D) Gene name prefixes enriched in cluster 3.3. Families shown have at least four members in cluster 3.3; proportion of the family located in this cluster is depicted in brown bars. Less than 10% of genes are in cluster 3.3 (“all”). (E) Gene prefix diversity (Shannon’s H) is lowest in cluster 3.3. (F) Distribution of promoter, intron, flanking sequence, and coding sequence GC% across genes which do not have paralogous neighbors in their home isochore (gray, genes with 0 tandem neighbors), genes which exist in small local duplications (red, genes with 1-4 tandem neighbors), and genes which exist in large multigene arrays (brown, genes with 5 or more tandem neighbors). A subset of genes without introns appear as 0’s.

The power of this approach is that it captures patterns of feature GC% in relation to one another: For most gene categories, a sharp rise in GC content marks the approach of the TSS, while the transcribed region and the region 3’ of the TES share lower GC content. This GC rise at the TSS clearly corresponds to the promoter. In this context, the paltry GC enrichment at the promoters of genes in cluster 3.3 (and to a lesser extent 3.1) is extremely stark (Figure 3A).

Based on high-confidence annotation of transcription start sites, we showed previously that mouse olfactory receptor promoters share this GC-poor pattern (13). At that time, the TSS’s of other highly tissue-specific genes had not been mapped. Current human annotations in the MANE set are high-confidence, curated gene models. Nevertheless, many of the genes in cluster 3.3 are extremely tissue-specific (see below) and have less supporting mRNA data than more widely expressed genes. We identified 240 genes in the MANE set (1.3%) that lack annotated 5’ UTRs, i.e. where the annotated transcription start site and translation start site are the same. While these were indeed mostly contained in cluster 3.3 (150 of 1636 genes in 3.3, 9%), there was no difference in promoter GC content between these suspect gene models and other genes in cluster 3.3 (data not shown). Indeed, our manual inspection of available RNA data for a subset of these genes suggests that the transcription start sites are correct, but that translation likely initiates at a downstream ATG.

We next examined how a gene’s isochore GC context relates to the sequence content of its promoter and coding region. We found that patterns of local GC content of genes predicted the GC content of their home isochore, consistent with the statistical genetic null hypothesis, but very surprising considering the functional implications (Figure 3B). The GC-content of a gene with its flanking regions (gene extent with 25kb on each side) correlated closely with the GC content of its whole isochore (Figure S3C). Individually, promoter and coding sequence GC% were also positively correlated with isochore sequence content, but the correlation coefficients were weaker: A subset of genes in AT-rich isochores have GC-rich promoters or GC-rich coding sequences (Figure S3D, E).

Genes in clusters 3.1 and 3.3, lacking GC enrichment in their promoters, were highly enriched for the same functional categories as were genes in AT-rich isochores: chemosensation, xenobiosis, and defense/barriers. Indeed, as can be seen in Figure 1A, B and Figure S2D, sometimes entire arrays were members of cluster 3.3 (brown color). To systematically test this, we plotted the promoter GC content distribution of genes in categories we term “outward-looking” (chemosensation, defense, xenobiosis, barriers) versus “inward-looking” (e.g. transcription, kinase function, morphogens). Outward-looking genes have AT-rich promoters while inward-looking genes have GC-rich or average promoters (Figure 3C). We manually annotated common prefixes and enrichment of genes in cluster 3.3 (Figure 3D): This group included all the chemosensory families, many sets of digestive and detoxifying enzymes, and several receptor arrays in the immune system and skin. It also included clustered protocadherins, which share transcriptional regulation patterns with chemosensors. In accordance with the preponderance of tandemly arrayed genes found in cluster 3.3, we found that genes in this cluster were housed in fewer unique isochores and had lower name diversity than those in the other k-means clusters (Figure 3E, S3C).

Finally, we asked whether being located near paralogues could predict local sequence features (Figure 3F). Indeed, genes near 1-4 neighbors with the same prefix had more AT-rich promoters than genes not located near paralogous genes, and genes with more than four same-prefix neighbors had AT-elevated promoters, coding regions, and flanking regions compared to both genes in small clusters and singletons. The striking coordination of isochore, promoter, and coding sequence AT content in tandemly arrayed “outward-looking” gene families prompted us to investigate how this pattern relates to the evolution of tandem arrays.

### Increasing AT content during evolutionary expansion of tandem arrays

As shown above, we find that both the regional and local AT content is high in tandemly arrayed gene clusters in human. AT bias could have emerged as gene families expanded or could have been pre-existing and supported molecular mechanisms of gene duplication. To ask whether AT content rises with the number of paralogous genes in a tandem array, we sought to track copy-number-variable tandem arrays across mammals. Assessing the evolution of tandem arrays is difficult, as large proportions of local paralogues can be species-specific duplications (Figure S4C and (55–58)). Therefore, instead of seeking to identify true homologous genes across different mammalian species, we used synteny analysis to follow whole arrays over evolutionary time. We searched for large gene arrays in human that could be identified across diverse mammals using microsynteny, where conserved heterologous genes serve as “bookends” bounding the ends of the paralgous tandem array.

While copy-number-invariable gene arrays, such as the HOX clusters, were easy to track across all mammals, as expected, the copy-number-variable gene arrays that would be informative for this analysis showed frequent assembly errors, micro-inversions, and array invasions by other heterologous genes. For example, while we observed that the expanded KLR array of NK cell receptors in the fruit bat *Rousettus aegyptiacus* and pheromone-associated MUP array in *Mus musculus* each had sharply higher AT content than surrounding genomic regions, each array had a large assembly gap (9,59). While this limited the number (and, indeed, the extremity) of arrays that we were able to track, we identified six copy-number-variable arrays that were suitable for analysis: An OR array adjacent to the hemoglobin beta genes (60); the Cyp2ABGFST cluster of xenobiotic enzymes; the SERPINA cluster of defensive protease inhibitors; the “epidermal differentiation complex” (EDC), containing skin proteins in the LCE, S100, and SPRR families; and two clusters of Keratin-associated proteins (KRTAPs) which form epidermal appendages such as hair, nails, and claws. These six clusters have diverse median GC content as can be seen in Figure 4A-H. We also included the copy-number-invariable HOXA cluster for comparison.

**Figure 4:**
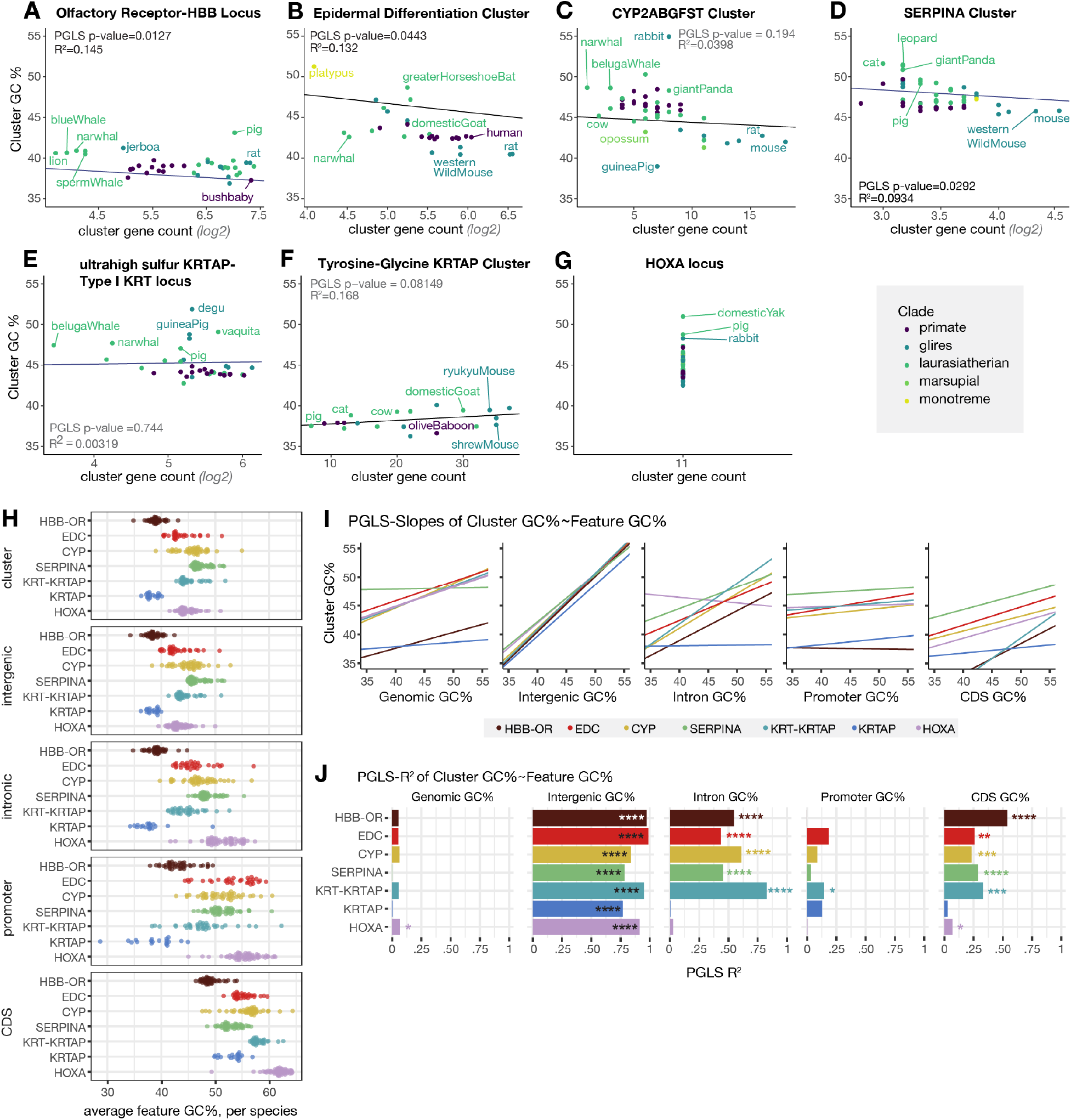
AT content is correlated with tandem array expansion. (A-G) Comparison of number of (A) OR genes in the Hemoglobin β (*HBB*) cluster, (B) LCE, SP100, and SRR genes in the Epidermal Differentiation cluster, (C) CYP2A genes in the CYP2A-GFST cluster, (D) SERPINA genes, (E) KRTAPs and KRTs in the Type I KRT locus, (F) KRTAPs in the high tyrosine-glycine KRTAP cluster, and (G) the HOXA locus with cluster GC% across mammals. All p-values report results of phylogenetic least squares analysis (PGLS). (H) Average GC% across the cluster, intergenic, intronic, promoter, and coding sequence of each tandem array. Each point represents one mammal species from panels A-G. (I) PGLS slope values and (J) R-squared values of phylogeny-corrected correlation of feature GC% to cluster GC%. Asterixes represent magnitude of PGLS p-values for each relationship, i.e. one asterixes represents a p-value between 0.05 and 0.01, two represent a p-value between 0.01 and 0.001.

For each of these clusters, we defined bookend genes as indicators of synteny that must be present for a particular species to be included in analysis. In mammals for which we could find these bookends on the same scaffold or chromosome, we computed the number of genes between bookends, spot-checking species whose counts were markedly higher than those of other species. The number of paralogues in each cluster is plotted across the mammalian clade in Figure S4C, showing the frequency of lineage-specific cluster expansions and contractions. Our full annotations of the contents of these clusters are provided in supplemental Table 6.

For each cluster in each species, we calculated the GC% between the bookend genes; relationships between GC% and gene count appeared to be log-linear. We performed phylogenetic least squares (PGLS) analysis to test for a relationship between GC content and log(paralog number) while controlling for phylogenetic relatedness in the multi-species dataset (Figure 4A-H). For the OR, EDC, and SERPINA clusters, AT content rose significantly with paralogue number (PGLS *P < 0.05*), suggesting that AT content can rise as the number of paralogs in tandem arrays grows. Certain arrays in certain species will have latent assembly or annotation errors; however, for the HBB-OR array, we manually counted a subset of species in parallel and saw the same trend (Figure S4A). The HOXA cluster showed variation in GC% despite maintaining the same number of paralogs in the cluster over evolutionary time; this variation appears to be predicted by evolutionary variation in the length of the cluster (i.e. number of base pairs between bookends, not shown). We were able to follow the HBB-OR cluster to non-mammalian amniotes, and found a sharp rise in GC content as the number of ORs in the region melted to 0 (Figure S4A, B).

The raw GC% of each cluster was poorly correlated with variation in genome-wide GC content across species (Figure 4I, J). However, each cluster type maintained a consistent relationship to the overall GC% of the genome: Across diverse mammalian species, the OR cluster and KRTAP cluster were almost always more AT-rich than the genome as a whole, while the other five clusters were almost always more GC-rich than the genome as a whole (not shown).

What portions of these genomic regions drive the variation in local GC content over evolutionary time? To assess this, we developed a series of scripts we call TandemClipR (insert github link once available), which divided each cluster in each species into CDS, intron, promoter, and intergenic regions. To avoid the “false-TSS” problem described above, we included only genes with annotated 5’ UTRs in our promoter analyses. We found that the GC% of each sub-region was positively correlated with the overall cluster GC% (Figure 4H-J). As we described above, this is expected from a statistical point of view, but surprising from a molecular point of view given that coding regions and promoters perform important molecular work. For example, only some amino acids can be coded with high-GC codons. Variation in intergenic regions contributed the most to cluster GC%, as expected based on their comprising a preponderance of the sequence. Despite varying with cluster GC%, coding regions, introns, and promoters had consistently higher GC content than the cluster as a whole, consistent with their functional constraints (Figure 4H).

Finally, we asked how the GC% of sub-regions of the cluster were related to the number of paralogues in the cluster (Figure S4D,E). For the clusters whose AT content rose with local paralogue number, the AT content of each substituent portion of the sequence (promoters, coding regions, introns, intergenic regions) also rose. The particular feature that correlated best with paralogue number varied across gene families, and no individual feature was consistently more correlated with paralog number than GC% of the whole cluster. In sum, we find that cluster AT content can rise as paralogous clusters bloom over evolutionary time; remarkably, these trends affect all the sequence components of the cluster, suggesting neutral or selective mechanisms that act cluster-wide. We hypothesize that local gene contents, particularly paralogous clusters of genes, can influence the emergence of isochores differing in GC content. Next, we provide a model for how this could have come about.

### Accumulation of Functional Diversity and Divergence

As we show above, many types of outward-facing genes have extreme variation in paralogue number across mammalian species (Figure S4C). Anecdotally, genes in these families also exhibit extreme allelic diversity and copy number variations among humans, and polymorphisms in these genes underlie human phenotypic variation in drug metabolism, sensory perception, and immune response (1,11,55,61–66). Colloquially, outward-looking genes are so diverse in copy number and sequence that a first step in GWAS is often to “throw out the ORs.”

Previous reports, including ours, have speculated that partitioning inward- and outward-looking genes into different parts of the genome could enable a higher ongoing mutation rate in AT-rich, outward-looking genes; however, point mutation rate is in general positively correlated with GC content because cytosines are especially mutation-prone (13,42,67,68). In the first model, AT content would facilitate mutagenesis and diversification of genes in these families. In the second model, relatively high point mutation drift in large gene families could have contributed to their shift to higher AT content over evolutionary time.

To test between these possibilities, we first systematically examined the degree of coding sequence variation in human genes grouped by AT/GC content or by degree of local tandem gene duplication, using the gnomAD dataset of rare single nucleotide variants ascertained from whole exome sequencing of >100,000 unrelated people (gnomAD v2.1.1, Figure 5A, B) (69). The ratio of protein-altering versus synonymous variants is positively correlated with AT content, with genes in AT-rich isochores and k-means cluster 3.3 highly enriched for potentially functional variants (Figure 5A,B). Moreover, genes near paralogous neighbors had higher levels of functional variants (Figure 5E,F). We note that use of rare variants profoundly understates the allelic variety in outward-looking genes, which exhibit radical common variation and high rates of copy number and structural variation. For example, any two humans are estimated to have function-changing variation in 30% of their olfactory receptor genes (70,71).

**Figure 5:**
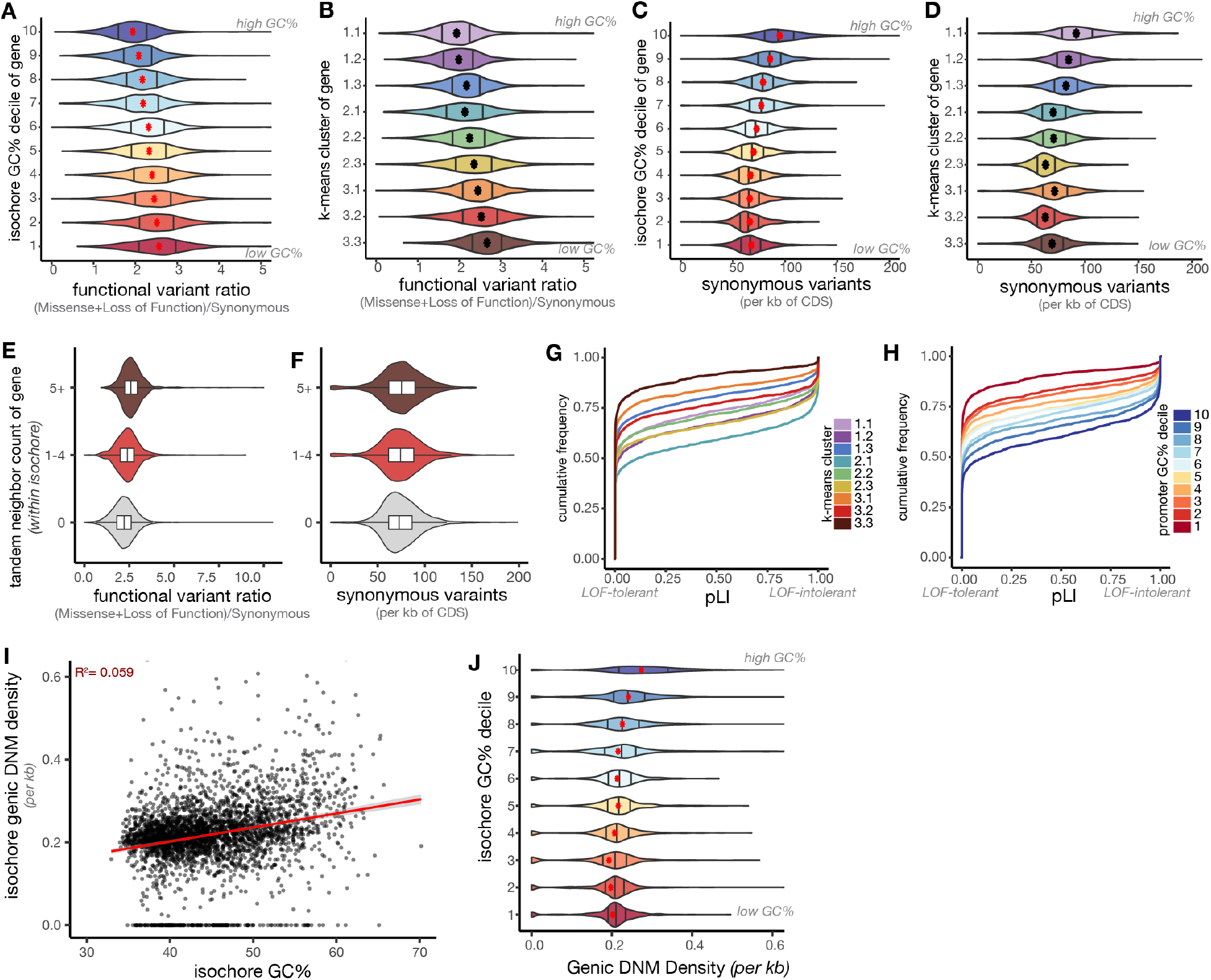
AT-rich genes have high functional diversity despite experiencing moderate mutation rates in the present. (A,B) Ratio of functional (missense plus loss of function) versus synonymous rare variants in the MANE gene set identified in gnomAD v2.1.1 exome sequencing of >100,000 unrelated individuals (69). Genes are binned by isochore decile (A) or k-means cluster (B), and dots indicate means. gnomAD rare variants are defined by < 0.1% allele frequency. (C, D) Raw counts of rare synonymous variants per gene in gnomAD v2.1.1 binned by isochore decile (C) or k-means cluster (D). (E,F) Functional (E) and synonymous (F) variant rates from gnomAD v2.1.1 for genes with or without paralogous neighbors. Box plots show median and mid-quartile distribution. (G,H) Cumulative frequency distribution plots of gnomAD pLI (likelihood that a gene is loss-of-function intolerant) relative to a gene’s k-means cluster assignment (G) or promoter GC% (H) (69). (I,J) Number of *de novo* point mutations observed per kb across the genes (TSS to TES) within an isochore relative to isochore GC% (I) and isochore GC% binned by decile (J). ∼700,000 DNM calls are pooled from all ∼11,000 trios sequenced to date (75).

Does the high polymorphism of genes in outward-looking tandem arrays result from differential mutation or selection versus inward-looking genes? To examine mutations directly, we used synonymous variants from gnomAD as a proxy. We see that AT-rich genes have *fewer* synonymous variants across unrelated people than do GC-rich genes (Figure 5C, D, S5A-D). This is consistent with point mutation rate being grossly driven by deamination of cytosine, especially in the C^me^G context: AT-biased genes have essentially run low on CpG dinucleotides to mutate (32–36,72,73). Therefore, we reject the hypothesis that AT-biased genes are “prioritized for mutation;” rather, they are enriched for functional diversity relative to a low ongoing mutation rate. We observe that olfactory receptor genes have higher rates of variant calls than do other AT-rich genes (Figure S5F). This inflated rate could reflect an unknown mutagenic process but could also result from incomplete knowledge of the full human “OR-ome” and incorrect assignment of homology relationships.

The enrichment of functional variation in outward-looking genes relative to low levels of synonymous variation could result from different selective processes acting on single-versus multi-copy genes. Single-copy genes are uniquely responsible for a particular biological process, while tandem arrays may distribute a particular process over a large set of family members (i.e. subfunctionalization). Deleterious mutations to single members of such large gene families are therefore more likely to be of scant phenotypic consequence. Indeed, we find that genes with more tandem neighbors have higher rates of function-changing versus synonymous variants (Figure 5E,F, S5E). To test how local and isochore GC content relate to the functional dispensability of genes, we used the gnomAD metric of “loss-of-function intolerance” (pLI): genes have high pLI if deleterious mutations are depleted from the population (69,74). Genes predicted to be loss-of-function intolerant were enriched in GC-rich isochores, had GC-rich promoters, and were absent from cluster 3.3 (Figure 5G,H, S5G); AT-skewed, outward-looking genes had lower probabilities of being loss-of-function intolerant, i.e. they are relatively dispensable.

To examine modern mutagenesis patterns in a different way, we sought to measure the rate of *de novo* mutations that occur in genes in AT-versus GC-biased regions of the human genome. Whole genome sequencing of two parents and a child (trios) enables detection of *de novo* mutations (DNMs). We used a recent dataset that compiles DNMs from >11,000 trios, nearly all those who have been sequenced to date (75). While these data remain sparse relative to the size of the genome (∼700,000 total DNMs), DNMs approximate a record of mutagenesis that has yet to be operated on by external selection. Pooling DNMs across each isochore and across transcriptional units (TSS to TES) within that isochore, we found that both genic and isochore-wide DNMs were more common in higher-GC isochores (Figure 5I,J, S5H-I). This is consistent with sequence-based predictions, prior findings, and our analysis of gnomAD synonymous variants within genes (76,77). Together, comparison of *de novo* mutations in meiosis and single nucleotide variants in unrelated humans both support the conclusion that despite their high divergence and diversity, AT-biased genes experience fewer contemporary mutations than do GC-biased genes, allowing us to reject the hypothesis that quarantining outward-looking genes in AT-rich regions of the genome facilitates higher mutation rates. Instead, lack of purifying selection on outward-looking gene blooms could be one factor that increases their AT content.

### Recombination and PRDM9 Binding

Recombination may be the primary influence on large-scale patterns of AT/GC content due to GC-biased gene conversion, which elevates GC content. AT-biased chromosomal bands are described as experiencing less recombination than GC-biased bands (20,26,30). We sought to test how recombination rates relate to GC content of inward-versus outward-looking genes. We therefore used whole genome sequencing data from human trios to measure recombination in isochores of different GC contents and in their constituent genes (78).

Crossovers appeared rare within gene blooms (Figure 6A, S6B). We calculated a relative crossover rate for each isochore and found that AT-rich isochores experienced less maternal and paternal crossovers than GC-rich isochores, as has been previously observed, though maternal crossovers were sharply diminished in the highest-GC isochores (Figure 6C,D) (15,78–80). We noticed that these low-crossover, high-GC isochores were often at chromosome ends, where maternal recombination has been shown to be low (81). To systematically examine recombination relative to each gene along the chromosome, we generated a Manhattan plot of crossover rate for each gene and its flanking regions (Figure 6B, S6A). This highlights the higher recombination of genes located in GC-rich isochores, except for maternal recombination at chromosome ends.

**Figure 6:**
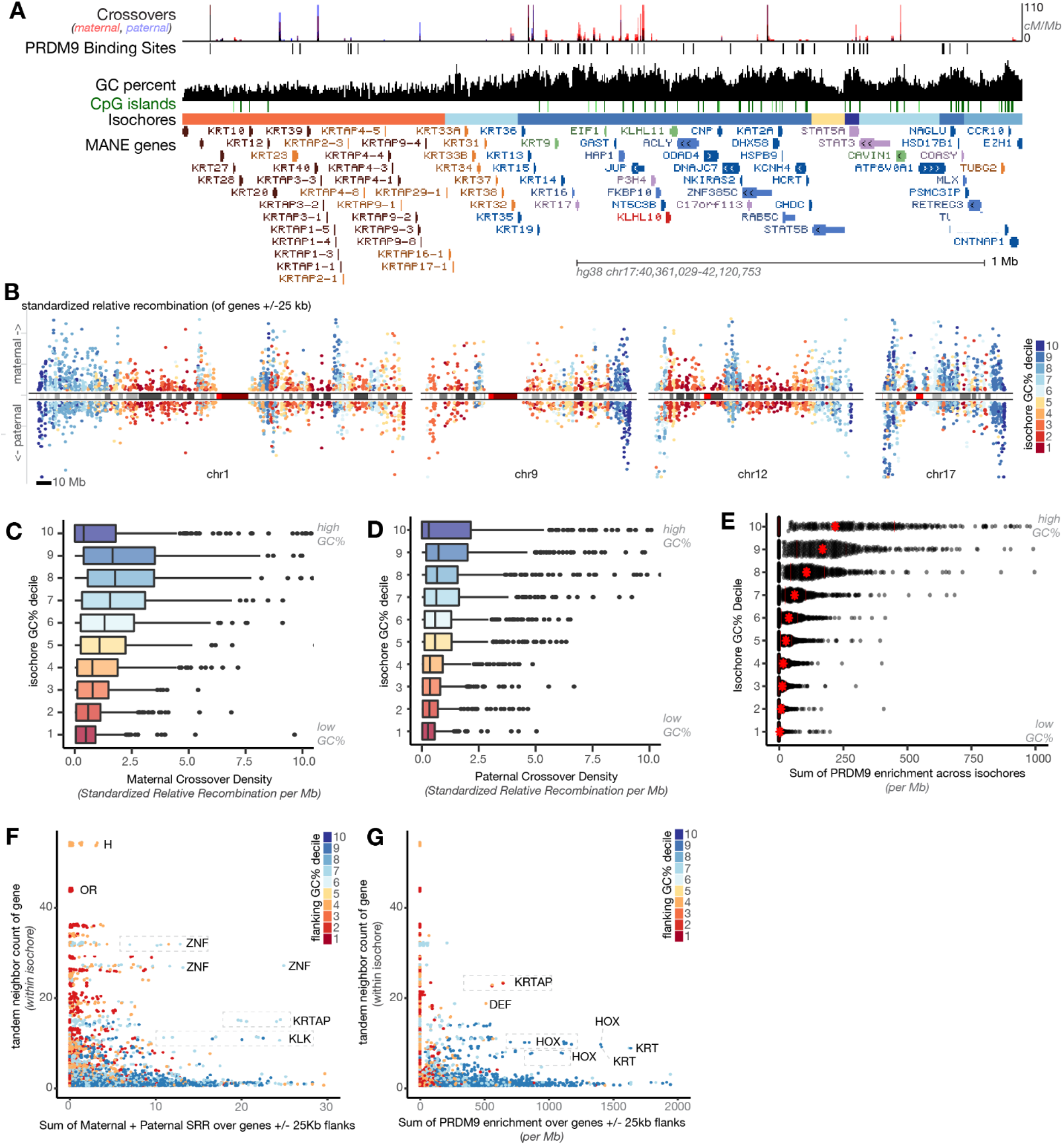
Crossovers are directed away from AT-rich isochores and genes. (A) UCSC genome browser screenshot of human KRTAP cluster showing recombination rates (maternal meiosis in red, paternal meiosis in blue) and PRDM9 binding calls (black). (B) Manhattan plot of standardized maternal (above 0) and paternal (below 0) recombination rate for each gene with its 25kb flanking regions. Genes are colored according to the GC content of their home isochore (Red: high AT. Blue: high GC). Scale bars: 10Mb. Chromosome ideograms reflect centromeres (bright red), gaps (dark red) and Giemsa bands (grays). Recombination rates are low for genes located in AT-rich isochores, except at chromosome ends, which are depleted for maternal recombination. (C) Maternal and (D) paternal crossover rates in isochores binned by GC percent. Crossover calls are from deCODE (78). (E) PRDM9 peak enrichment across isochores binned by GC%. (F) Scatterplot of tandem neighbor counts per genes within the same isochore versus (F) the sum of maternal and paternal standardized relative recombination and (G) sum PRDM9 enrichment.

Recombination in paralogous clusters can induce dramatic insertions, deletions, and chromosome rearrangements if paralogues errantly pair with one another (82). While clusters of outward-looking paralogues are enriched in AT-rich regions, the conserved clusters of paralogues located in GC-rich regions, such as Hox clusters, may also be risky to recombine. To separate the influence of AT/GC content versus clustered-ness, we compared recombination rates across clusters of different sizes (Figure 6F). However, most large clusters of genes were AT-rich and had little recombination, making it difficult for us to separate which variable dominates. While recombination is generally directed away from genes, some genes experienced crossovers. We plotted the isochore and k-means cluster distribution of genes with the 10% highest internal crossover rate (TSS-TES, Figure S6D, E). These genes were in GC-rich isochores and excluded from AT-rich k-means cluster 3.3. Together, these analyses suggest that gene blooms located in AT-rich regions of the genome experience low current crossover rates.

In many vertebrates, including humans, crossovers are seeded by binding of PRDM9 to its target site (83–85). To test whether variation in observed recombination across AT/GC categories is due to differential seeding, we examined PRDM9 binding data from human cells (Figure 6E,G, S6F,G)(86). We observed a striking depletion of PRDM9 binding from AT-biased gene clusters and isochores, in line with previous analyses (79). We note that many animals have lost PRDM9, and in the absence of PRDM9, recombination is often seeded at CpG islands (87). As described below, AT-biased gene families also lack CpG islands. Taken together, these results suggest that AT-biased, tandemly arrayed gene clusters experience low rates of recombination, and that this likely results from low rates of crossover initiation.

### Transcriptional regulation of outward- versus inward-looking genes

Our analyses above lead to the hypothesis that clusters of paralogues that are not subject to purifying selection (i.e. “outward-looking genes”) lose GC content as they bloom due to reduced recombination and increased tolerance of point mutations. We further hypothesize that the sequence content and context of these genes has been evolutionarily co-opted to produce extreme tissue-specificity and/or variegation of their transcription. While we expect that isochore-level AT/GC content influences genomic organization and histone mark flavors, which would influence transcription indirectly, we also noticed that paralogous clusters of genes lacked annotated CpG islands in their promoters (see CpG Island track, Figure 2A, B, Figure S2D)(54). CpG dinucleotides are depleted from vertebrate genomes due to the mutability of methylated cytosine; nevertheless, CpGs are relatively enriched in vertebrate promoters, and these “CpG islands” frequently remain unmethylated (88). Previous analyses suggested that 50-70% of mammalian genes have CpG island promoters (19,89,90). Nevertheless, by considering each gene in its sequence context, we estimate that 90% of protein-coding genes have GC enrichment directly upstream of the TSS (Figure 3A).

While CpG islands are calculated relative to the GC content of the region, we sought to ask if it is even *possible* to have a CpG island in an AT-rich region. As in Figure 2, we separated isochores of at least five genes by GC content, gene number, and prefix diversity, and then calculated CpG island scores on a per-gene basis (Figure 7A,B). Surprisingly, genes located in all types of isochores *could* have CpG islands in their promoters, but genes in large paralogous arrays often did not. Highly conserved genes in arrays, like Hox and Histone arrays, retained their islands regardless of the GC content of the isochores. Gene blooms lacking CpG islands tended to be outward-looking, and almost all large, island-poor arrays were in AT-rich isochores. We also noticed that the shape of islands around the TSS varied across our k-means clusters, with some clusters having islands that were symmetrical relative to the TSS, and others having islands that were polarized to the 3’ side of the TSS (Figure S7A).

**Figure 7:**
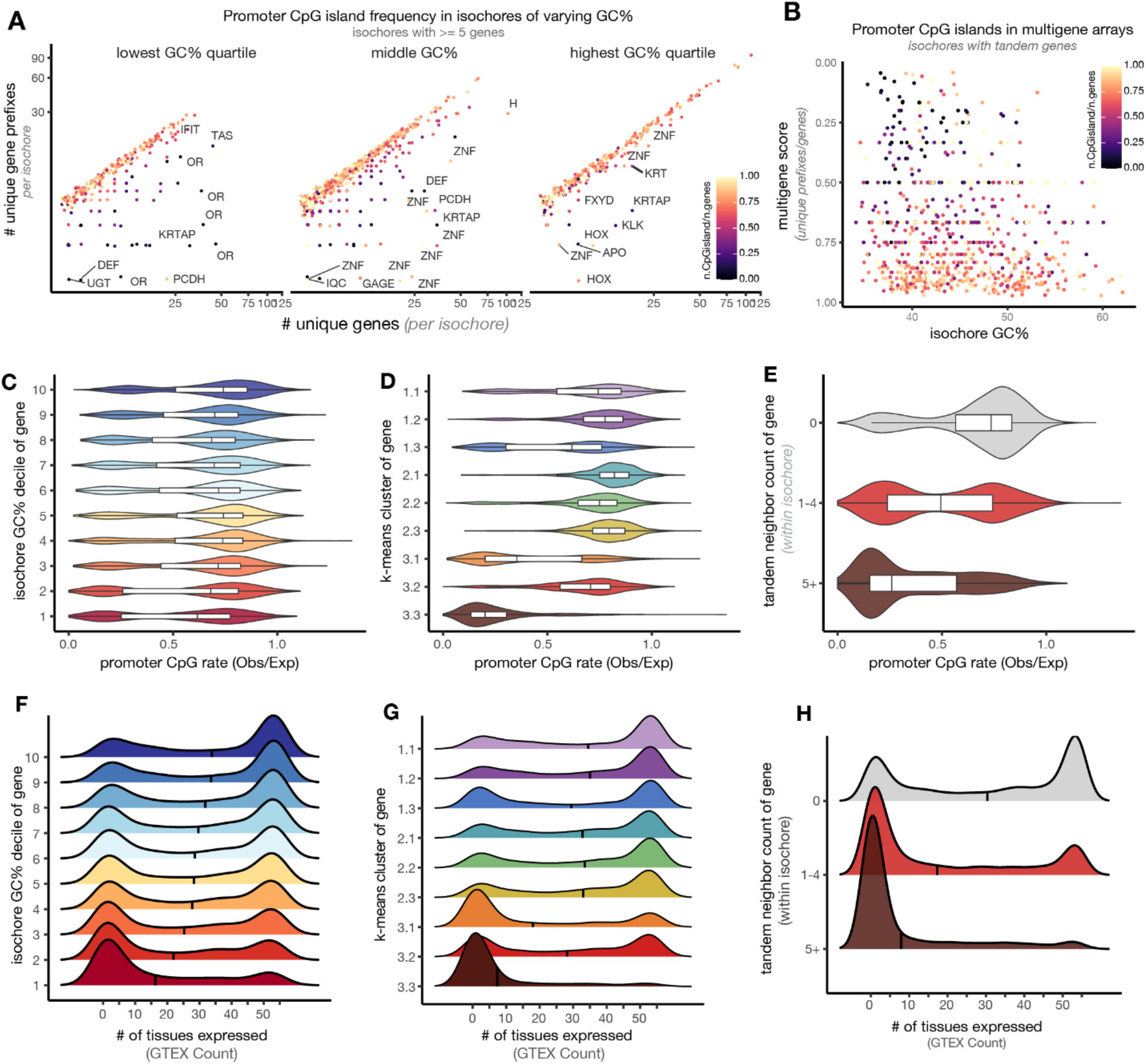
Lack of CpG islands and restricted expression is a feature of tandemly arrayed AT-rich genes. (A) Promoter CpG island frequency (from (54)) in isochores of varying gene content. Same plots as in Figure 1E, recolored to highlight promoter characteristics. Most common gene name prefixes in the isochore are labeled. (B) Multigene score versus isochore GC% for isochores containing tandem duplicated genes. Color represents fraction of genes in the isochore with promoter CpG islands. (C-E) Distribution of promoter CpG rate for genes binned by (C) isochore GC% decile, (D) k-means cluster assignment, or (E) tandem duplication score. We counted CpG’s from −750 to +250bp around the gene TSS and compared to the expected rate for a sequence of that GC content. (F-H) Distribution of tissue-level gene expression for genes binned by (F) isochore GC% decile, (G) their k-means cluster assignment, or (H) tandem duplication score. GTEx data from 54 human tissues was binarized to “expression” or “no expression” based on RPKM of 5. AT-rich genes and those in paralogous arrays are detected in few or no sampled GTEx tissues, while GC-rich genes are frequently detected in all sampled tissues. Black bars depict medians.

Because CpGs are only protected from methylation when they are located near other CpG’s, i.e. in islands, and because methylated CpG’s are extremely mutation-prone, CpG loss can be precipitous once it begins (32). We calculated observed promoter CpG sites compared to the expected rate based on local sequence content and observed a bimodal distribution across genes of all isochores, though the observed/expected CpG’s elsewhere in the isochore were predicted by GC content (Figure 7C,D, Figure S7A,B). Thus, though CpG-less promoters are more common in high-AT isochores, islands are compatible with genes in any isochore context; island loss appears specific to paralogous gene blooms (Figure 7E). Across a range of isochore GC contents, genes near tandem neighbors had CpG-poor promoters compared to singletons (Figure S7E). This promoter state also drives k-means clustering of genes in 3.1 and 3.3 (Figure 7D). Many genes in 3.1 and 3.3 have no more CpG’s in their promoters than the genome-wide average (∼25% of expected CpGs). Further separating these k-means clustered genes into those with paralogous neighbors and those without did not stratify promoter architecture, suggesting that the k-means categories are already sensitive to the AT/GC sequence features delineating genes in paralogous clusters (Figure S7D).

As we and others suggested previously in the mouse, CpG-less promoters are likely to be regulated by non-canonical mechanisms that are independent of TATA Binding Protein (TBP) (13,91). This could allow the unique and rare expression of these genes relative to their CpG-containing brethren: their ground state is to be “off forever.” While hemoglobin beta and amylase genes, which we now know to be small clusters of paralogues, were shown to lack CpG islands in their promoters in the 1980’s, measurements of the relationships between tissue-specificity and promoter architecture were performed before the transcription start sites and promoters of tissue-specific genes were mapped (15,19,92). To test how AT/GC distribution in genes relates to patterns of gene expression, we used GTEx data, which measures gene expression in 54 tissue types taken from adult human donor cadavers (93). We set a threshold (RPKM of 5) to binarize this quantitative data to “expression” or “no expression.” In agreement with Bird’s hypothesis, this simple metric varies sharply across genes of different k-means clusters or with differing promoter or home isochore GC content (Figure 7F-H)(13,15,19,92).

Genes that are GC-rich are often expressed in many or most tissues tested, while AT-biased genes most often appear to be expressed nowhere or in one tissue. We note the many genes with “no” expression in GTEx are specific to tissues not sampled by GTEx (e.g., olfactory receptor genes in the olfactory epithelium). We infer that genes not detected in any tissue in GTEx data—i.e. the preponderance of AT-rich genes—are highly tissue-, cell type-, time-, or condition-dependent in their expression. As we have shown throughout this study, these exotically transcribed, CpG-less genes are located in evolutionary dynamic clusters of paralogues and tend to be housed in AT-rich isochores.

In the longest-lived cells in the body, post-mitotic neurons, tandemly arrayed gene families have been shown to be clustered with one another in nuclear space and to be uniquely protected from accumulation of CpH (Cp-nonG) methylation (47,94,95). These findings, together with the overwhelming transcriptional repression of these gene families, suggest that they might be sequestered in nuclear space away from the transcriptional machinery. Indeed, we found that across 21 tissues sampled by Hi-C, AT-rich genes and genes located in AT-rich isochores were likely to be located in transcription-suppressing “B” compartments (Figure S7F, G)(96). Previous analyses have demonstrated that variation in GC content also predicts local chromatin looping and association with the lamina and other nuclear structures (16,39,49,97).

## Discussion

Animals make extensive and varied contacts with the external environment, both engaging with foreign molecules and producing and excreting their own substances. Specialization of these input-output functions plays a definitive role in animal lifestyle, and often occurs in mammals via amplification and diversification of tandemly arrayed gene families (10,12,13,98,99). Extensive gene losses are also common—for example, just as the vomeronasal organ is vestigial in humans, human genes for vomeronasal receptors are no longer functional (100,101). Here, we define a common genomic architecture in mammals for genes whose products engage the external world: they are in tandem arrays, are found in AT-biased isochores, and lack CpG islands in their promoters (Figure 1) (13,44). In humans today, we find that genes in AT-rich tandem arrays experience low rates of point mutation and recombination, and tend toward tissue-specific expression patterns.

### Modeling the emergence of AT/GC sequence bias in outward-looking tandem arrays

The rapid turnover of genes in tandem arrays by birth-and-death evolution can complicate efforts to reconstruct their evolutionary history. Using synteny, we were able to track paralogue number changes for a set of polymorphic arrays in mammals (Figure 4). For arrays of olfactory receptors, skin proteins, and defensive protease inhibitors, AT content increases with copy number across mammals. This analysis suggests that high AT content need not be an ancestral feature of gene blooms, but can emerge during array expansion. Using population genetic data from humans, we test whether elevated rates of allelic diversity in outward-looking genes results from distinct mutational or selective effects. We accept the argument of Lynch that AT/GC content itself is unlikely to be directly operated on by selection, and instead focus on neutral and selective events affecting the genes contained in sequence-biased ischores (27).

We find that genes in AT-biased tandem arrays experience low ongoing rates of point mutation (Figure 5) and low rates of recombination (Figure 6). We suggest that excessive allelic diversity in these regions could have arisen due to weakened selection on historical point mutations. Because the most common point mutations replace C or G bases with T or A bases, tolerance of point mutations in expanding tandem arrays could shift GC content down; low rates of point mutation in the present would be due to the scarcity of mutable CpG dinucleotides remaining in these clusters (38). Tandem arrays also appear depleted for recombination events that can shift GC content back up (Figure 6). Together, tolerance of historical point mutation and strengthened intolerance of recombination as gene families expand would tend to bias expanding arrays toward higher AT content (Figure 1).

While deamination of cytosine in the CpG context dominates mutation rates, many other neutral processes contribute to the overall mutation spectrum, and these too operate differently in copy-number-variable tandem arrays. For example, AT-biased regions of the genome are late-replicating, which is associated with higher rates of germline mutations, and genes in tandem arrays tend not to be transcribed in the testis, which would exempt them from transcription-coupled repair (42,102). Excess allelic variation could also result from paralogous gene conversion. While paralogous gene conversion can be measured for paralogous pairs, in extended arrays this phenomenon would be difficult to distinguish from copy number variation without long-read data spanning the region (103). Similarly, genes in tandem arrays do not have one-to-one homology relationships across species. Trivially, this means that their interspecific divergence is high, but they would be left out of gene-wise measures of divergence that rely on clear homology assignments.

Other mutagenic and selective forces may also contribute to the evolution of AT-sequence bias. For example, the repetitive nature of these regions of the genome predisposes them to problems, such as replication fork slippage, that require non-homologous end joining (NHEJ) to resolve (104,105). RCC4-DNA ligase IV is primary ligase complex for NHEJ, and it preferentially ligates poly-dT single-stranded DNA and long dT overhangs (106). Polymerase mu, which is also involved in NHEJ, shares a similar bias towards both dT and dC, though preferentially towards dT (107). These biases could encourage these regions of the genome to adopting AT-rich sequences with each duplication, and thus each addition of repetitive material.

### Implications for gene regulation

Genes located in AT-biased tandem arrays are typically silenced almost everywhere in the body and expressed at extremely high levels at a specific place and time. Where and when these genes are expressed, they perform the definitive work of the cell type. Many of these outward-looking gene families exhibit some kind of exclusive expression, from the fetal to adult switch in hemoglobin β expression (i.e. exclusion over time) to the one-receptor-per-neuron pattern of olfactory receptor expression (i.e. exclusion over space). We expect that genes located in high-AT isochores that lack CpG islands in their promoters are durably silenced by particular mechanisms when they are not expressed (i.e. almost everywhere), and that their expression will be activated by non-canonical mechanisms in the single condition where each is expressed. As we and others suggested previously for mouse ORs, CpG-less promoters are likely to be regulated by non-canonical mechanisms that are independent of TATA Binding Protein (TBP) and the recently discovered basal CpG-island-binding factors BANP and BEND3 (13,91,108,109). As Bird hypothesized presciently in 1986, “When activated, they appear to be bound to a complex of tissue specific factors which presumably accomplish what…islands can achieve using ubiquitous, non-tissue specific factors. Given that there are usually rather few CpGs near tissue specific genes…one would not expect to find CpG or methylation built into the activation mechanism of genes of this type. (92)” One could argue that these genes do not have promoters at all and rely completely on locus control regions to concentrate and deliver transcription factors to the TSS (45,110,111). This atypical architecture appears to predispose these genes to restricted expression relative to their CpG-containing brethren: their ground state is to be “off forever.”

Many or most molecular genetic events are sensitive to variation in AT/GC distribution: AT content predicts compartmentalization of the genome in 3D space, replication timing, and patterns of histone marks (41,112–116). Typically, AT-biased sequence is packaged as heterochromatin and silenced. Work on olfactory receptors, clustered protocadherins, and secreted liver proteins suggest that these gene families are expressed from the context of constitutive heterochromatin, which appears to be present prior to expression and to be retained on family members that are not expressed (117–121). CpG islands also function as molecular beacons: they mark transcription start sites, serve as recombination hotspots in the absence of PRDM9, and act as replication origins in meiosis (87,112,122). The accrual of high AT content in gene arrays and the lack of CpG islands is therefore likely to exert a strong effect on the molecular regulation of these genes. In ectodermal development, single-copy genes accrue CpH methylation, perhaps passively, while AT-rich gene arrays remain devoid of this modification (94). This suggests that AT-rich gene arrays are locked away from the ambient molecular stew of the nucleus, perhaps over very long developmental time periods. Indeed, tandem arrays clump together in the nucleus in post-mitotic neurons (47,95).

In addition to the extreme time and/or tissue specificity of most outward-looking gene families, a fraction of these families exhibit stochastic expression such that each cell expresses just one or a sparse subset of family members. Chemosensors, B- and T- cell receptors, NK cell receptors, and clustered protocadherins all exhibit this restricted expression (119,123,124). As we argued recently, sparse cell-wise expression patterns compartmentalize the effects of mutations (119). These mechanisms are also likely to result in insensitivity to copy number variation, as each cell chooses its own dose of family members for expression. Feedback mechanisms that ensure cells can “choose again” if they originally pick a pseudogene further buffer potential deleterious effects of mutations (125,126). Combined with the sheer numbers of family members that a particular gene function is distributed across, these expression mechanisms could predispose these genes to selective drift.

### Role for recombination in diversification versus maintenance of gene arrays

Local sequence diversity and recombination rates are usually positively correlated (reviewed in (127)). In contrast, we find that AT-biased gene families exhibit high diversity and divergence despite low recombination. Comparison of the human and chimpanzee genomes found two distinct types of high-divergence regions: Genome-wide, chromosome ends are GC-rich and have high divergence and recombination; in chromosome interiors, however, recombination is more modest overall, and divergence is higher in G-bands, which are AT-rich (20). As Holmquist argued previously based on much more limited data, we show here that these G-bands contain very different types of genes than do Q-bands (15,48).

Overall GC content is theorized to be a record of past gene conversion, with high-GC regions having experienced high historical recombination (31). If this is the case, then AT-bias in tandem arrays could reflect historical depletion of recombination in these regions, while also dampening recombination rates in the present (Figure 6). Modeling suggests that once variation in AT/GC content starts to emerge, it can be self-reinforcing via positive feedback (32). There remains conflict between the mode by which these gene arrays are thought to have bloomed—i.e. via gene duplication through ectopic exchange during recombination—and their current depletion for recombination events. Other modes of duplication, including replication slippage and transposition, may also be at work in expanding these arrays, and duplication mechanisms could themselves influence GC content (128).

Ectopic exchange in repetitive gene regions can have benign or catastrophic consequences. Induction of copy number variation within a gene array may be of small phenotypic consequence, as the jobs these genes perform are by nature distributed across many family members. In contrast, ectopic exchange that deletes a cluster or induces recombination between clusters can result in catastrophic chromosome rearrangements (82). Retrotransposition or ectopic exchange mediated by repetitive elements can seed new gene clusters elsewhere in the genome (129,130), and gene families with more than one cluster genome-wide are likely to be particularly dangerous for genome stability. Indeed, mammalian chromosome evolution appears to have been shaped by ectopic exchange between OR clusters, to the extent that ORs are often positioned near chromosome ends (131–139). Finally, even if structural variation in outward-looking tandem arrays is benign within an individual, it can lead to hybrid incompatibility and can initiate or reinforce reproductive isolation that leads to speciation (140–142). Recent modeling work has sought to characterize the tradeoffs between the structural fragility of gene blooms and the potential positive effects of allelic diversification (143).

Given the genomic danger of these tandem arrays, why have gene family members remained *in cis* with one another? An extreme example is the “milk and teeth” locus on human chromosome 4. The casein genes in this locus evolved via tandem duplication of enamel genes at the root of the mammalian tree; the enamel genes themselves evolved from *follicular dendritic cell secreted protein* in bony fish (12,144). Astonishingly, these genes have remained syntenic. Why would this be the case, given that they’re expressed in three separate body systems and that such tandem arrays are genomically dangerous? We propose that as in the case of maintenance of Hox gene synteny, the regulatory elements of these genes remain tangled with one another, such that relocation of array members elsewhere in the genome would divorce them from cis-regulatory elements that they depend on for expression (145–147). Recent research on enhancer evolution in animals suggests that enhancer tangling can result in the preservation of synteny over ∼700 million years (148). In other cases, as in B Cell Receptor, hemoglobin, clustered protocadherin, interferon, and chemosensor arrays, family members share and compete for the same regulatory elements (111,111,149–152). This mutual dependence would again increase the phenotypic consequences of recombination events that break synteny by separating genes in large families from locus control regions they depend on for expression.

Array incompatibility between individuals of a species and the necessity of remaining co-located with regulatory elements that may be tangled with or shared by other gene family members could cause tandem arrays to behave like supergenes—multigene regions inherited as an allelic unit. We suspect that depletion of CpG islands and PRDM9 sites from tandemly arrayed genes protects the genome from the danger of errantly recombining these duplicative regions. Nevertheless, recombination and gene duplication or deletion still sometimes occur in these regions—their crossover rate even today is non-zero—and the marginal fitness effects of resulting copy number variants could allow products of these meiosis to be preserved in the population. As gene arrays get larger, point mutation tolerance could shift their GC content downward, putting the brakes on recombination as they become ever more unwieldy. Overall, recombination in these regions would be dampened, while differential tolerance of local duplications versus gross rearrangements could allow an increase in local allelic diversity.

### Implications for chromosome organization

Repetitive elements have shaped chromosomal evolution since the dawn of eukaryotes. The linear genome is proposed to have arisen from erroneous meiotic recombination between Group II introns which invaded the circular genome to create the t-loop precursors to stable telomeres (153). Similarly, dispersion and expansion of ORs and other large tandem gene arrays have shaped mammalian chromosome evolution. Tandem arrays of ORs represent ancestral breakpoints of chromosomal synteny between mice, rats, and humans (138,154). A large OR cluster is found at the end of the q-arm of chr1 in humans but not in mice. In addition to ORs, large gene families including zinc finger (*ZNF*) and immunoglobulin heavy chain (*IGH*) genes are observed at chromosome ends across eukaryotes (155). In the modern human population, unequal crossovers between OR clusters are a source of recurrent and pathological rearrangement hotspots (156).

While we also observe these AT-rich isochores *near* chromosome ends, terminal isochores are often some of the most GC-rich in the genome and can contain heterogeneous single-copy genes (157). Indeed, the largest OR cluster at the end of the q-arm of chromosome 1 in humans is followed by a higher GC% isochore containing ZNF genes. This strong end-GC% accumulation appears to arise paternally: genes in high GC% isochores at chromosome ends are enriched for paternal crossovers and relatively depleted of maternal crossovers. Overall, paternal crossovers are biased towards chromosome ends (81,158). Chromatin organization of pachytene spermatocytes is implicated in this phenomenon. Spermatocytes have shorter synaptonemal complexes compared to oocytes, and, futher, subtelomeric regions do not require PRDM9 for crossovers, however, how these factors contribute to paternal crossover end-bias remains incompletely understood (159,160). Potentially, recombination-based alternative lengthening of telomeres (ALT) in spermatocytes biases hotspots towards chromosome ends (161).

Over evolutionary time, as ectopic recombination places high-AT tandem arrays at chromosome ends, high paternal rates of gBGC at the ends of chromosomes would generate new isochores of increasing GC% comprising newly evolving genes (162,163).

### Is mutation biased or random with respect to gene function?

Recent mutation accumulation studies have suggested that *de novo* mutations could occur with different frequencies in different kinds of genes or in genic versus non-genic locations (164). We and others also argued previously that segregation of mutation-tolerant versus mutation-intolerant genes into AT-versus GC-biased regions of the genome could allow differential mutation rates on different classes of genes (13,67,68). However, our analyses of synonymous versus functional variant rate and of *de novo* mutation rate in AT-versus GC-biased genes suggest the opposite: that AT-biased genes experience fewer mutations in living humans than do GC-biased genes. This is in line with neutral, sequence-based expectations (42). While CpG prevalence dominates the mutation spectrum, other mutational and repair processes that differentially affect AT-versus GC-biased regions of the genome almost certainly contribute to the overall pattern of diversity in inward-versus outward-looking genes. In particular, AT-rich regions of the genome are late-replicating, and tandemly arrayed genes are excluded from transcription-coupled repair during spermatogenesis. Each of these patterns could increase the mutation rate in outward-looking genes. Late replication can worsen the loss of cytosines, while lack of TCR would be expected to increase the rate of A>G (T>C) mutations (42,102).

While mutagenic or repair processes specific to gene blooms could contribute to their overall mutation spectrum, we expect that for most, allelic diversity and differential AT/GC content in inward-versus outward-looking genes result from differential selection trajectory over evolutionary time. One attractive model is that as a gene cluster expands in size and the function of that gene family is partitioned over more and more members, purifying selection becomes weaker and weaker on individual family members (Figure 1). Because the most common point mutations are C->T and especially CpG->TpG, evolutionary tolerance of point mutations would shift GC content down. On the other hand, purifying selection and higher recombination rate would both preserve the GC content of singleton genes.

### Matching sequence content for genes and their isochore environments

Throughout this study, we show that the local AT/GC content of genes correlates closely with that of their isochore environment (e.g. Figure 2B, S2B-G). We found this pattern perplexing. For example, while higher purifying selection on single-copy genes can help to explain their higher GC content, we would expect most variants in the non-coding portions of an isochore to be phenotypically neutral. Because gene conversion acts over longer chromosomal distances, differential recombination could affect the sequence content of genes and their environment together. This pattern could also result from background/linked selection, which is known to vary in strength across the genome (165,166). Because function-changing variants are physically linked to local variants that don’t alter phenotype, differential purifying selection on mutations in single-copy versus multi-copy genes would affect the strength of background selection in the neighborhood and inheritance of linked neutral variants. Background selection will operate over shorter distances in regions with higher recombination (i.e. GC-rich isochores), but because multiple inward-looking, single-copy genes are co-located in the same isochore, there may be little nearby that is exempt from linked selection. In this way, purifying selection that results in maintenance of high GC content in single-copy genes would also preserve the high GC content of their isochore neighborhood. Finally, variation in repetitive element distribution could also contribute to the AT/GC content of isochores housing outward-looking versus inward-looking genes. Indeed, increased LINE density in chemosensory gene families has been proposed to contribute to their regulation (167).

The matching sequence content of genes and their isochore neighborhoods would seem to facilitate coherent patterns of histone marks across entire isochore units and their organization in nuclear space. On the other hand, genes with AT-versus GC-rich coding sequences use distinct codons, which could affect their translation rates given variation in the prevalence of various tRNAs. Further analysis will be required to assess whether amino acid distribution varies for proteins coded by AT-rich versus GC-rich sequences.

Finally, while we can find genes with GC-rich promoters in all kinds of isochores (Figure S3F), we never find genes with AT-rich promoters in GC-rich isochores. While we favor the model shown in Figure 1, where isochore-level AT content rises during gene blooms due to the parallel actions of protein coding drift and suppressed recombination, we cannot exclude (and are indeed fascinated by) the possibility that the distinct regulatory characteristics that allow LCR-mediated expression could be the keystone factor tying outward-looking genes to AT-rich isochores. However, in the absence of other evidence, we think that LCR-driven transcription is a kludge that allows expression of paralogous clusters that failed to maintain their CpG’s, rather than CpG-less promoters emerging via positive selection to allow regulation by LCRs.

### Is this genomic architecture specific to mammals?

While isochore structure is not unique to mammals, it is not a universal feature across animal clades, and the AT/GC variation observed in mammals is extreme (27). We are curious whether stem mammals evolved molecular mechanisms that facilitated the evolution of gene arrays. These could include both systems that maintain these arrays as constitutive heterochromatin when they are not being expressed and unique transcriptional mechanisms that activate them, often in a stochastic or highly restricted manner, in their target tissues. One candidate factor that mediates long-range enhancer-promoter interactions in multiple arrayed families is Ldb1 (45,168). Social insects have also massively expanded their olfactory receptor gene repertoire in cis; in the ant, this is accompanied by increased AT content (169). Have convergent mechanisms for stochastic expression facilitated olfactory receptor repertoire expansion in insects?

Other clades may have evolved distinct mechanisms to organize repetitive genes or gene pieces: In *Diptera*, repetitive arrays are often organized as alternative splicing hubs (170–173). Reptiles and birds exhibit “microchromosomes” which have distinct GC content from the rest of the genome and can house arrays of rapidly evolving, outward-looking genes such as venoms (174). Trypanosome arrays of surface VSGs are located in subtelomeric regions (175). For mammals, the “isochore solution” balances diversity in gene arrays with genomic integrity.

## Author Contributions

Conceptualization, MVB, MAC, EJC; Methodology, MVB, MAC, MLH, EJC; Investigation, MVB, MAC, MLH, EJC; Formal Analysis, MVB, MAC, MLH; Visualization, MVB; Data curation, MAC; Writing – Original Draft, MVB, MAC, EJC; Writing – Review & Editing, MVB, MAC, MLH, EJC; Funding Acquisition, EJC; Supervision, EJC.

## Acknowledgements

Thanks to Rachel Duffié, Sebastian Zöllner, David Lyons, and members of the Clowney lab for discussion and comments on the manuscript. This work was supported by the Rita Allen Foundation Milton Cassel Scholarship and the Alfred P Sloan Research Scholarship in Neuroscience to EJC. EJC is a McKnight Scholar and a Pew Biomedical Scholar. MVB was supported by the NIH Cellular and Molecular Biology Training Grant T32-GM007315. MAC was supported by the NIH Early Stage Training in the Neurosciences Training Grant T32-NS076401.

## Descriptions of supplemental files

**Supplemental Table 1:** Isochore calls in the hg38 human genome assembly.

**Supplemental Table 2:** Isochore calls in the hg19 human genome assembly.

**Supplemental Table 3:** Results of all statistical analyses presented here.

**Supplemental Table 4:** Genewise data, including isochore assignment, k-means cluster, local sequence characteristics, variant rates.

**Supplemental Table 5:** Description of data in Supplemental Table 4.

**Supplemental Table 6:** GC contents of selected gene families and their annotated features (ie: CDS, introns, intergenic regions).

**Supplemental Table 7:** List of genomes searched with Tandem ClipR.

## Methods

### Describing isochores

To call isochores, we implemented a genomic segmentation algorithm called GC-Profile (50) using halting parameter (number of segmentation iterations) of *t0* 275 and minimum segment length of 3000 bp. Gaps less than 1% of the input sequence were filtered out, generating 4328 distinct isochores in hg38 (Supplemental Table 1). Isochores were ranked by average GC%, with rank 1 having the highest and 4328 having the lowest. We also performed this analysis in hg19 (Supplemental Table 2). Isochores are reported in Supplemental Tables 1-2.

### Statistical analyses

We performed Kruskal-Wallis non-parametric ANOVA for each group of comparisons (Supplemental Table 3). We then used Dunn’s pairwise analysis to compare individual groups with one another (Supplemental Tables 3). We use phylogenetic generalized least squares (PGLS) regression as implemented in caper (176) to test for correlations between genomic characteristics across vertebrate species while controlling for the evolutionary non-independence of the multi-species dataset. We used timetree.org (177) to produce our species phylogeny for these analyses, where the tree was imported and pruned to the subset of species for a given comparison using the ape package (178). In PGLS, lambda was optimized by 0 and 1 by maximum likelihood.

### GC content calculations

Genes from the Matched Annotation dataset from the NCBI and EMBL-EBI (MANE) Select dataset (51) were downloaded from the UCSC Genome Browser. Isochores were matched to genes using the coordinates of the transcription start site. GC content across gene features, including promoters (−750 to +250bp flanking TSS), flanking regions (+/- 25kb), coding exons, exons and UTRs, and introns were separately calculated from FASTA sequences using bedTools (179).

To generate 9 k-means clustesr, we used gc5BaseBw from the UCSC Genome Browser (53) to calculate GC% scores across MANE genes with +/- 1kb flanks. We generated 3-kmeans clusters of genes, which were further clustered into 3-kmeans clusters each using deepTools plotHeatmap (180). Cluster assignment and quantification of other parameters for each gene are reported in Supplemental Table 4.

### Characterizing types of genes

To characterize the types of genes residing in isochores of varying GC, we used 2 categories of descriptors: GO terms and gene prefixes. To identify GO terms associated with genes in each isochore GC decile, we used the R package, clusterProfiler (version 4.2.2)(181). This helped us streamline identification of key terms that appeared in each decile. With this list, we identified GO terms that were most significantly enriched in each decile with a depth of at least 30 genes. Using AmiGO (182), an online database of GO identifiers, we pulled the list of genes associated with our selected group of significant GO terms and plotted GC content across each term. The terms we chose are listed in the table below.

| Shortened Term (from Fig 2) | Full GO Term Description | GO ID |
| --- | --- | --- |
| wnt signaling | wnt signaling pathway | GO:0016055 |
| kinase activity | kinase activity | GO:0016301 |
| transcription | transcription, DNA-templated | GO:0006351 |
| defense | immune response | GO:0006955 |
| xenobiosis | xenobiotic metabolic process | GO:0006805 |
| keratinization | keratinization | GO:0031424 |
| chemosensation | detection of a chemical stimulus | GO:0009593 |

We wanted an alternative to GO analysis for assessing diversity across isochores and k-means gene clusters. Since the prefixes of well-annotated genes (like the ones from the MANE dataset) are shared across genes within the same gene family, we used this as a means of assessing diversity with more specificity than one would achieve through GO analysis. The process of assigning gene prefixes is as follows:

1. Convert old names into new nomenclature.

- Go to the HUGO Gene Nomenclature Committee’s (HGNC)(183) website and the list of gene symbols from the MANE set into their “Multi-symbol checker” (the link provided will take you there directly). This will ensure we have the most up-to-date names for each of our genes (ie: some which may have been labeled as ‘FAM’ may have a new symbol to go with the rest of the gene family).
- Match names in the MANE set to names labeled “Approved symbols” by HGNC, and replace those symbols with the HUGO names.
2. Replace numbers with “_”. We can’t remove all numbers because there are several genes that have more letters after numbers that aren’t important for our purposes (ie: CSN2A will become CSN_A).
3. Remove anything after the first instance of “_” (ie: CSN_A will become CSN). The goal of this step is to keep the first part of the prefix, removing letters and numbers that indicate subfamilies.
4. While it isn’t common, some genes require us to know those numbers to know what they do (most commonly, enzymes involved in modifying carbohydrates). Largely, these genes start with a single letter, followed by numbers, then more letters. Thus, to fix these genes, we pull out the genes that have 1 letter after steps 4 and 5.
5. Look through each of those genes that start with only one letter, then decide how best to group them.
6. View gene prefixes in alphabetical order and search for prefixes that are likely to be families, then rename (ie: KCNT and KCNQ are both potassium channels, so we grouped these together).

Once we had a list of gene prefixes, we calculated a Shannon’s H diversity metric for each isochore based on the prefix probabilities in each isochore (proportions + log2(1/proportions) = diversity metric). Larger values are indicative of more diversity. Similarly, we calculated a Shannon’s H diversity metric for each k-means cluster.

### De novo mutations

*De novo* mutations (DNMs) were compiled by (75) from seven family-based whole genome sequencing (WGS) datasets, encompassing a total of 679,547 single nucleotide variants (SNVs), which comprise data from both neurotypical and neurodivergent individuals. We remapped the dataset to hg38 using LiftOver in UCSC Genome Browser. To calculate genomic DNM density, we counted the number of DNMs occurring within the coordinates listed in the GC calculation section above. To calculate DNM density, we pooled genic DNMs within each isochore and divided by the sum of the region of interest’s size, i.e. we identified all the genes in an isochore, summed the DNMs between their transcription start and termination sites, then divided by the summed length of those genic regions.

### Allelic variants

We used the gnomAD v2.1.1 dataset of single nucleotide allelic variants (69). The authors defined rare single nucleotide variants (<0.1% allele frequency) from 125,748 exomes and 15,708 whole genomes and predicted whether variants within coding regions are likely to be functionally synonymous, missense, or loss-of-function. Here, we used observed synonymous, missense, and loss-of-function mutation rates. We ported variant calls to MANE genes in hg38 using the gene symbol and Ensembl transcript IDs. In Figure 5, we also use the calculated pLI score from gnomAD, which describes the likelihood that a gene is loss-of-function intolerant in humans.

### Recombination

Crossover data for hg38 was acquired from deCODE where the authors used whole-genome sequence (WGS) of trios and were able to refine crossover boundaries for 247,942 crossovers in 9423 paternal meioses and 514,039 crossovers in 11,750 maternal meioses (78). Of note, the data we used here is restricted to autosomes. To calculate crossover density, we assigned crossovers to a region of interest based on the median of the crossover coordinates. We normalized counts within a region by dividing by the genomic average for that sex. PRDM9 binding data from HEK293T cells transfected with the PRDM9 reference allele was acquired from (86). We selected the top 10% of PRDM9 peaks based on enrichment scores to account for weak PRDM9 binding sites associated with overexpression in the system, as noted by the authors. Like with crossovers, we calculated PRDM9 binding site density across genes as the summed enrichment scores across genic regions within an isochore, mapping by the midpoint of the binding coordinates.

### Gene Regulatory Information

To determine tissue specificity of gene expression, RNA-sequencing data was sourced from Genotype-Tissue Expression (GTEx) project (V8, released in August 2019), containing 17,382 samples collected from 54 tissues from 948 donors (93). For each gene in the MANE set, we counted the number of tissues in which expression was at least 5 transcripts per million (TPM).

To measure A/B compartment occupancy of genes across tissues, AB compartments were sourced from published Hi-C data from 21 tissues and cell types (96). MANE genes were lifted over into hg19 to match A/B compartment domain calls in hg19. Isochores called in hg19 were assigned to a compartment by matching the isochore’s midpoint to the midpoint of the closest compartment. Genes were assigned to a compartment by matching the transcription start site to the midpoint of the nearest compartment, as most genes did not fall into a single compartment (∼90%). Further, we counted the occurrences of compartments A and B for each isochore and gene. These counts were binned into always A (21 counts of A), mostly A (14-20 counts of A or 0-6 counts of B), equally A or B (7-13 counts of A or B), mostly B (0-6 counts of A or 14-20 counts of B), and always B (21 counts of B).

To identify genes with CpG islands in promoter regions, we downloaded the CpG Island track (unmasked) from the UCSC Genome Browser (54). The ratio of observed vs. expected CpG dinucleotides was converted to a bigwig coverage file and plotted across gene TSS’s (+/- 2.5kb) in 9 k-means clusters using deepTools plotHeatmap. The average score of CpG islands within −750bp and +250 bp of a gene TSS were calculated using bedTools.

### Gene Bloom Evolution with Tandem ClipR

To access and analyze syntenic tandem gene blooms across species, we created a pipeline we call TandemClipR that defines tandem arrays based on orthologous “bookend” genes. Bookends define syntenic array bounding positions and are not members of the bloomed family. First, we used the biomaRt R package (184) to define the human genome as the reference mart, and then created a table of bookending ortholog chromosome names, starts, and stops using the ‘getLDS’ function and the dec2021 ensembl archive as a source. We tabulated these ortholog positions in all 193 vertebrate species in the archive (See Supplemental Tables 6 and 7). If both orthologs were on the same scaffold, indicating a contiguous assembly of the tandem array, we used the retrieved coordinates to subset the gff3 annotation file to the focal region using the subsetByOverlaps() function in the GenomicRanges package (185). Filters were applied when imported annotations to retain only “gene”, “mRNA”,“CDS”,“exon”,“five_prime_UTR”, and “three_prime_UTR” annotations.

The resulting gff3 files were used to analyze feature GC content across each tandem array based on sequences in the Ensembl “toplevel” assemblies. Namely, we used command-line tools to subset the gff3 annotations into focal features (genes, exons, five_prime_UTR), and used while awk and bedtools merge and subtract (179) were used to produced genic, intergenic, exonic, intronic, and whole cluster bedfiles defining these regions. Additionally, bedfiles defining the promoter region of each gene were produced by defining a window 750bp upstream and 250bp downstream of the transcription start site. We fed these bedfiles to bedtools nuc to calculate GC content for each feature, which were counted and aggregated into average GC% for final analyses. We spot-checked gene counts in NCBI’s Genome Data Viewer and noticed that there were a few cases in which counts varied between recent assemblies. For example, the lion HBB-OR cluster is much shorter in the assembly we used compared to the most recent assembly. However, our manual counts of the HBB-OR array in a subset of species produced similar trends as the automated annotation. To further verify that one of these assembly/annotation errors did not sway the overall trend, we ran each PGLS analysis iteratively, removing one species at a time, and found that PGLS p-values of the whole were largely reflected in each iteration.

---

**Figure S2.**
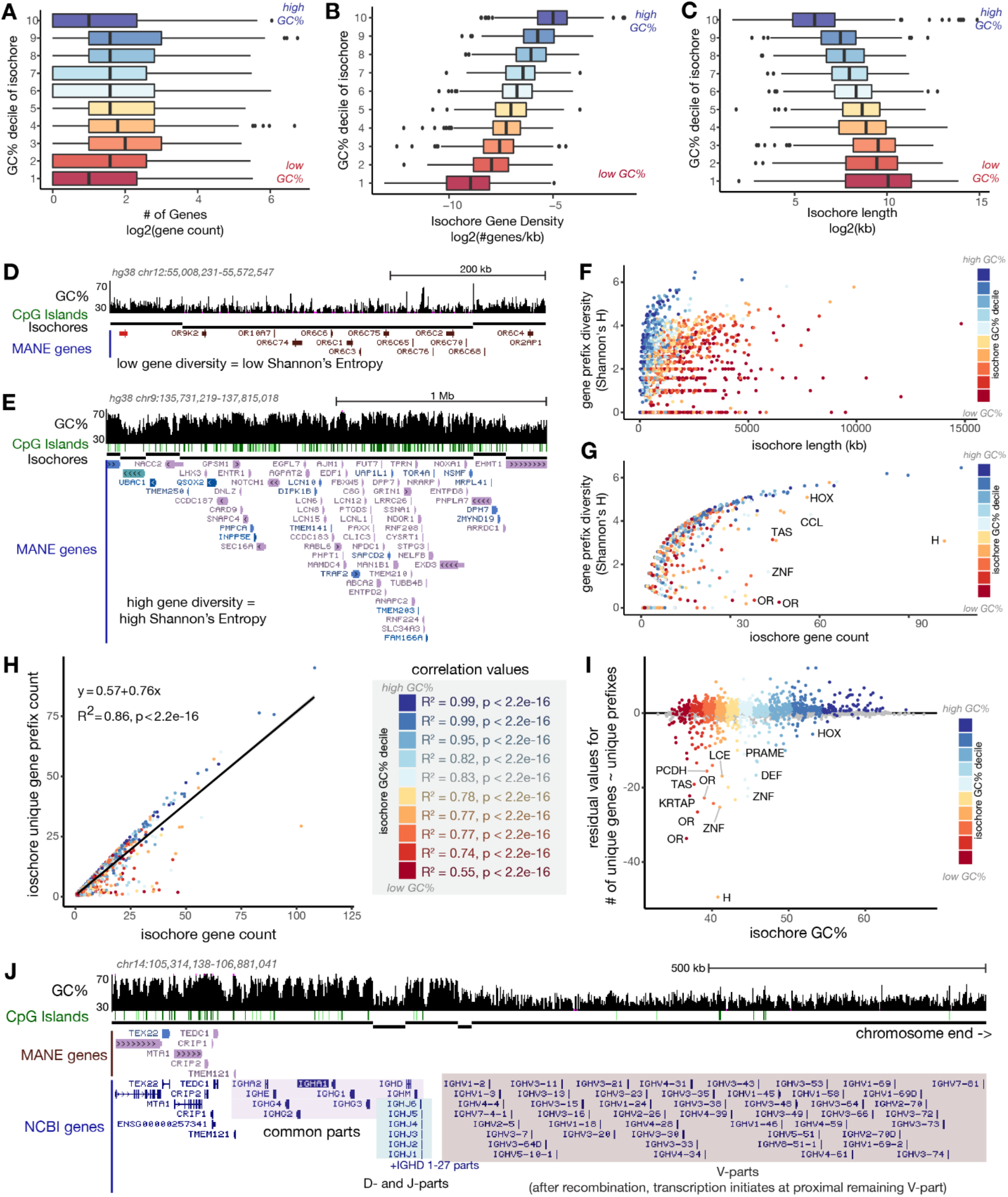
Characterization of human genomic isochores. (A-C) Box and whisker plots describe the relationship between isochore GC% and gene number (A), gene density (B) and isochore length (C). Isochores are binned into deciles according to GC content with decile 10 representing high GC and decile 1 representing high AT. (D,E) UCSC Genome Browser screenshots showing additional example isochores. Gene models are colored according to k-means clusters described below. (D) This isochore is AT-rich and contains a gene bloom; all the genes in this isochore have the same prefix, so prefix diversity (Shannon’s H) is low. (E) This isochore is GC-rich and contains genes from a variety of families, so diversity is high. (F,G) Relationship between isochore length (F) and gene count (G) and gene prefix diversity. Isochores are labeled by most common occurring gene prefix. (H) Full graph in Figure 2E including all gene-containing isochores. Spearmann correlation values per isochore deciles are shown on the right. (I) Residuals (i.e. distance of each point from regression line) of graphs in Figure 2E, S2H. (J) The *IGH* locus, containing V, D, J, and common regions of human IgH. V, D, and J regions are classified as “gene parts” and are not represented in the MANE set; however, the V repeats, where transcription initiates, are in a single AT-rich isochore and lack CpG islands.

**Figure S3:**
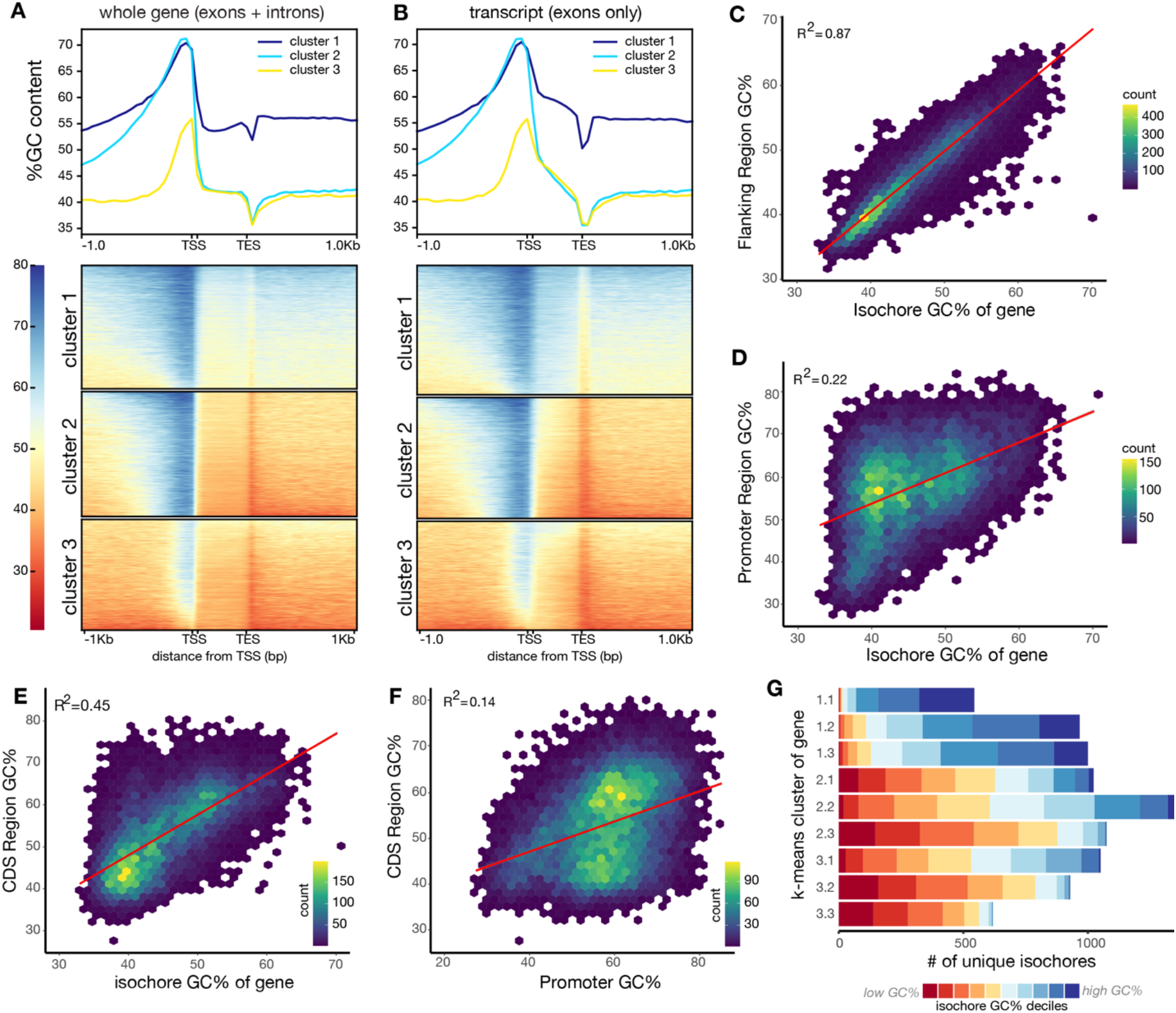
Relationship between GC content of local gene features and the home isochore. (A-B) Heatmaps displaying GC% over 50bp sliding windows calculated across genes. Graphs in (A) include introns, graphs in (B) exclude introns. Clusters were determined by k-means. Summary line plots depict mean GC% across clusters. <10 genes switched clusters when introns were excluded. (C-F) 2D histograms depicting the relationship between (C) flanking, (D) promoter, (E) coding region, and GC percent of “home” isochore for each gene in the MANE set. (F) Relationship between promoter GC percent and coding sequence GC percent. “Flanking region” spans from 25kb upstream of TSS to 25kb downstream of TES. Correlation coefficient (R^2^) and trend (red line) are shown. (G) Number of unique isochores housing genes in each k-means cluster. Genes in the more extreme k-means clusters are contributed by fewer isochores.

**Figure S4:**
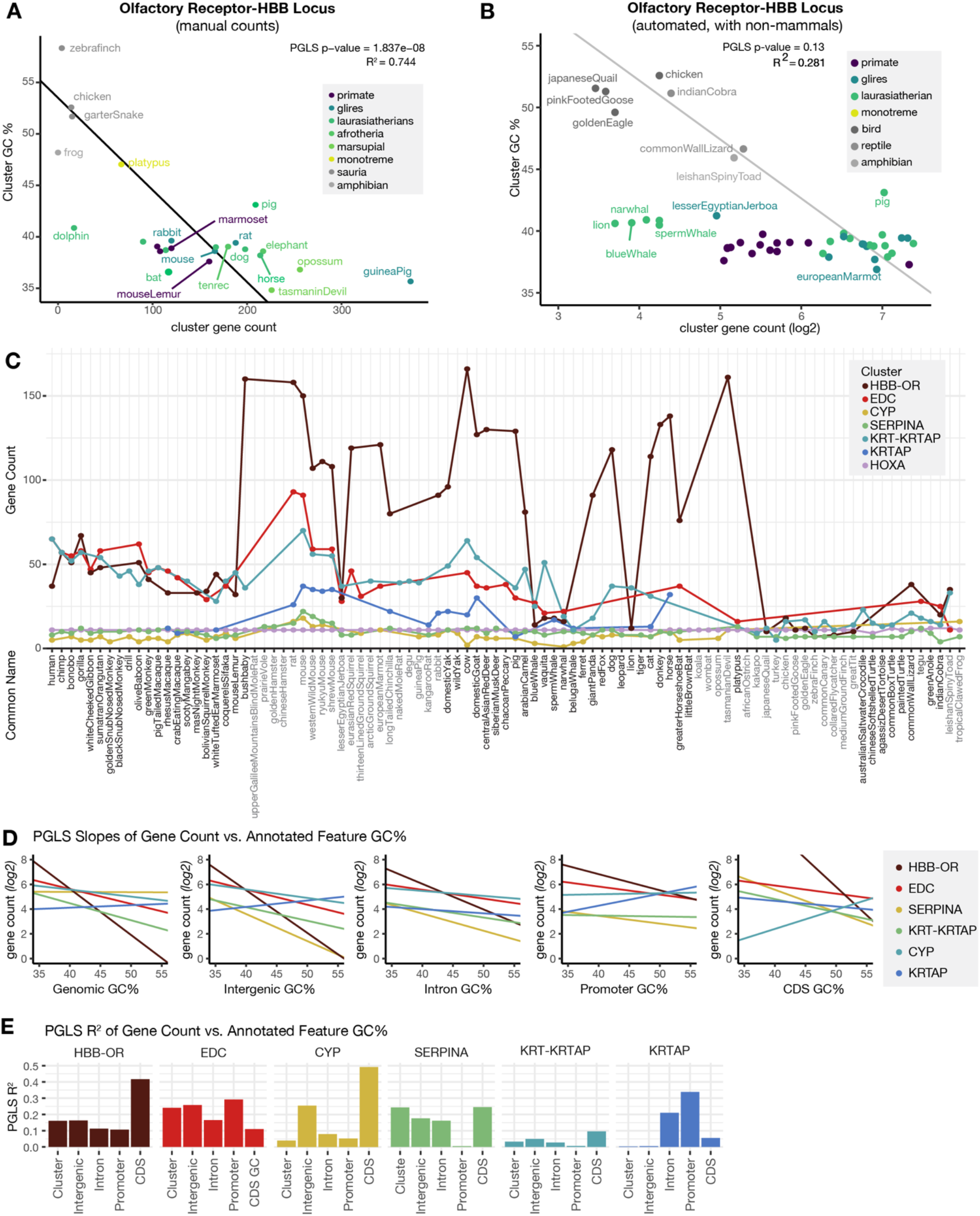
Data related to Figure 4. (A) Counts of gene number for each paralogous cluster across amniotes analyzed (B) UCSC Genome Browser screenshot showing the hemoglobin β cluster on human chromosome 11. As described previously, the *HBB* cluster is flanked by olfactory receptor genes and these are bracketed by conserved single-copy genes, including *STIM1*, *RRM1* and *FHIP1B* shown here, that are syntenic with the *HBB* cluster since before mammals branched (60). (C) Analysis of gene number and GC% in the HBB-OR cluster across mammals, reptiles, and birds. (D) R^2 values for the relationships between the GC% of each cluster and the GC% of its constituent parts across mammalian species. (E) R^2 values for the relationship between cluster or constituent part GC and the number of genes in the cluster across mammalian species.

**Figure S5:**
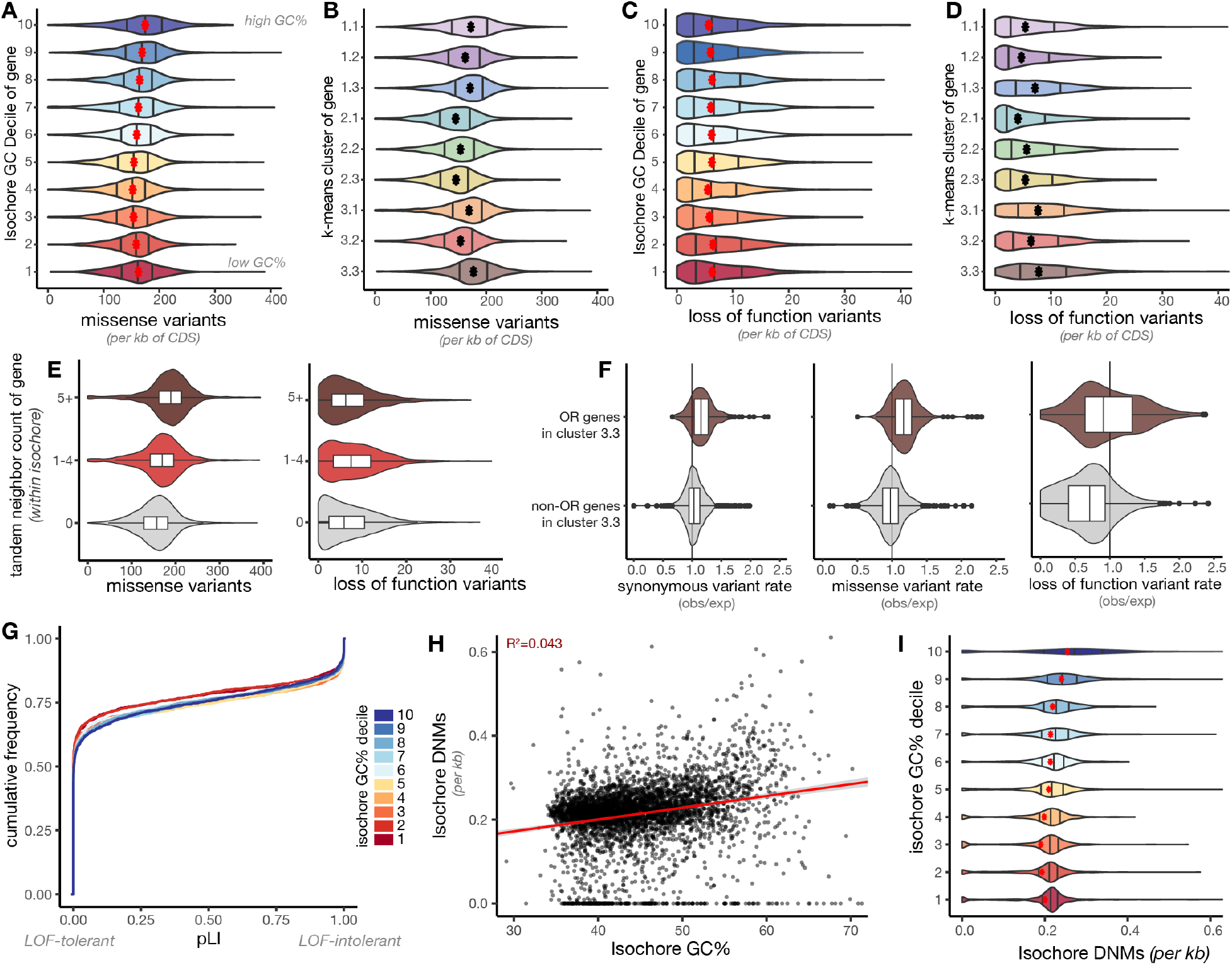
Raw rates of loss of function and missense variation. Raw gnomAD v2.1.1 counts of missense (A, B) and loss-of-function (C, D) SNVs across MANE genes binned by isochore GC% decile (A, C) or k-means cluster (B, D). (E) Missense and loss of function mutation accumulation in genes of differing degrees of local tandem duplication. (F) Ratio of observed over expected rates of synonymous variants, missense variants, and loss of function variants of genes in k-means cluster 3.3, split by OR and non-OR genes. Observed/expected values are calculated relative to the neutral mutation spectrum by gnomAD. (G) Cumulative frequency of pLI scores for genes binned by their home isochore GC% decile. (H,I) Number of *de novo* point mutations observed per kb in each isochore plotted relative to isochore GC% (H) and isochore GC% binned by decile (I). ∼700,000 DNM calls are pooled from all ∼11,000 trios sequenced to date (75).

**Figure S6:**
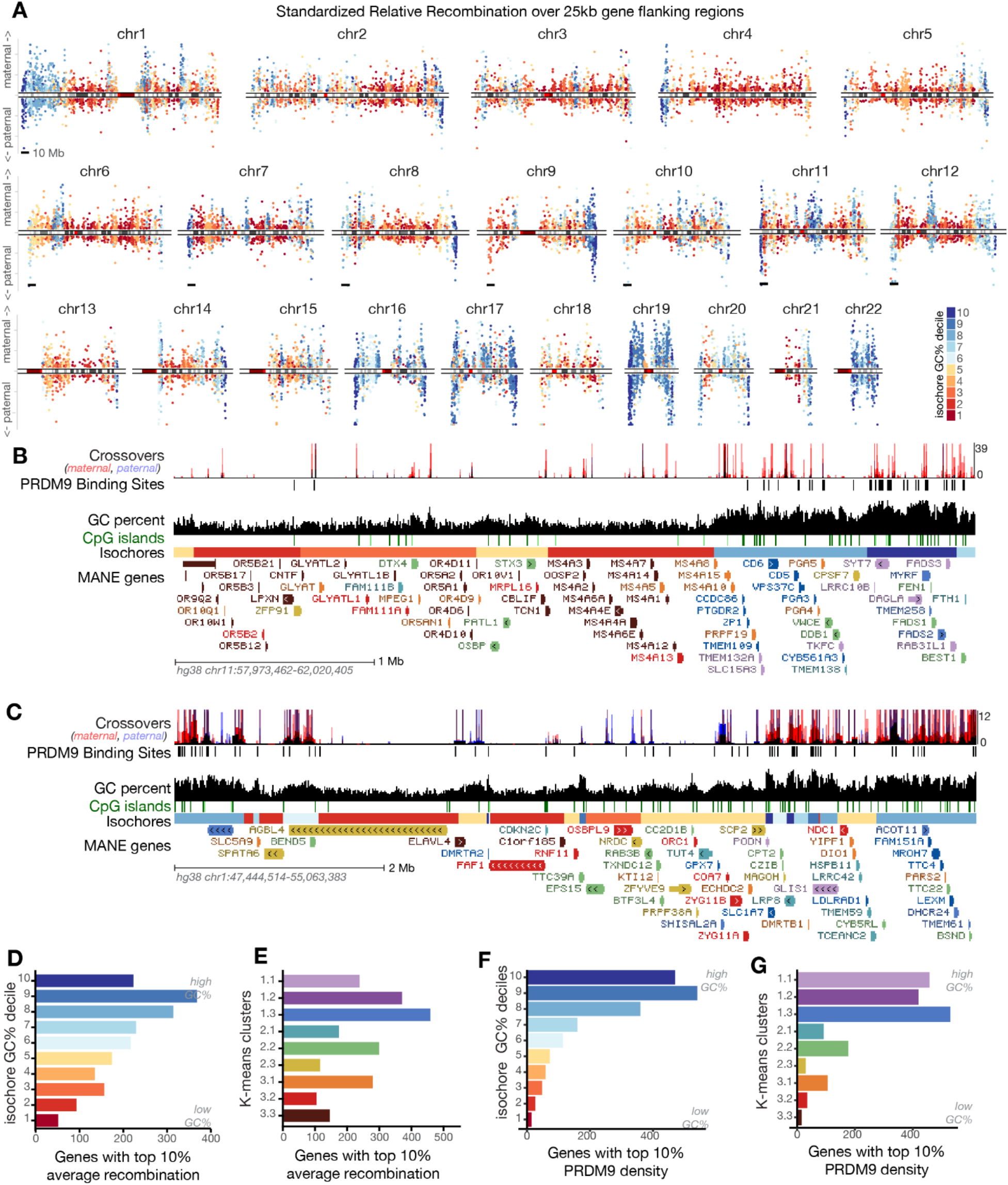
(A) Manhattan plots of maternal and paternal standardized relative recombination rates for all chromosomes, as described in Figure 6B. (B-C) UCSC Genome Browser screenshots of (B) OR and MS4A loci, flanked by GC-rich isochores and (C) interspersed GC and AT-rich isochores. (D,E) Counts of (D) isochore decile and (E) k-means cluster distribution of genes with 10% highest rate of maternal and paternal within-gene crossovers. (F,G) Counts of (F) isochore decile and (G) k-means cluster distribution of genes with 10% highest rate of within-gene PRDM9 binding.

**Figure S7:**
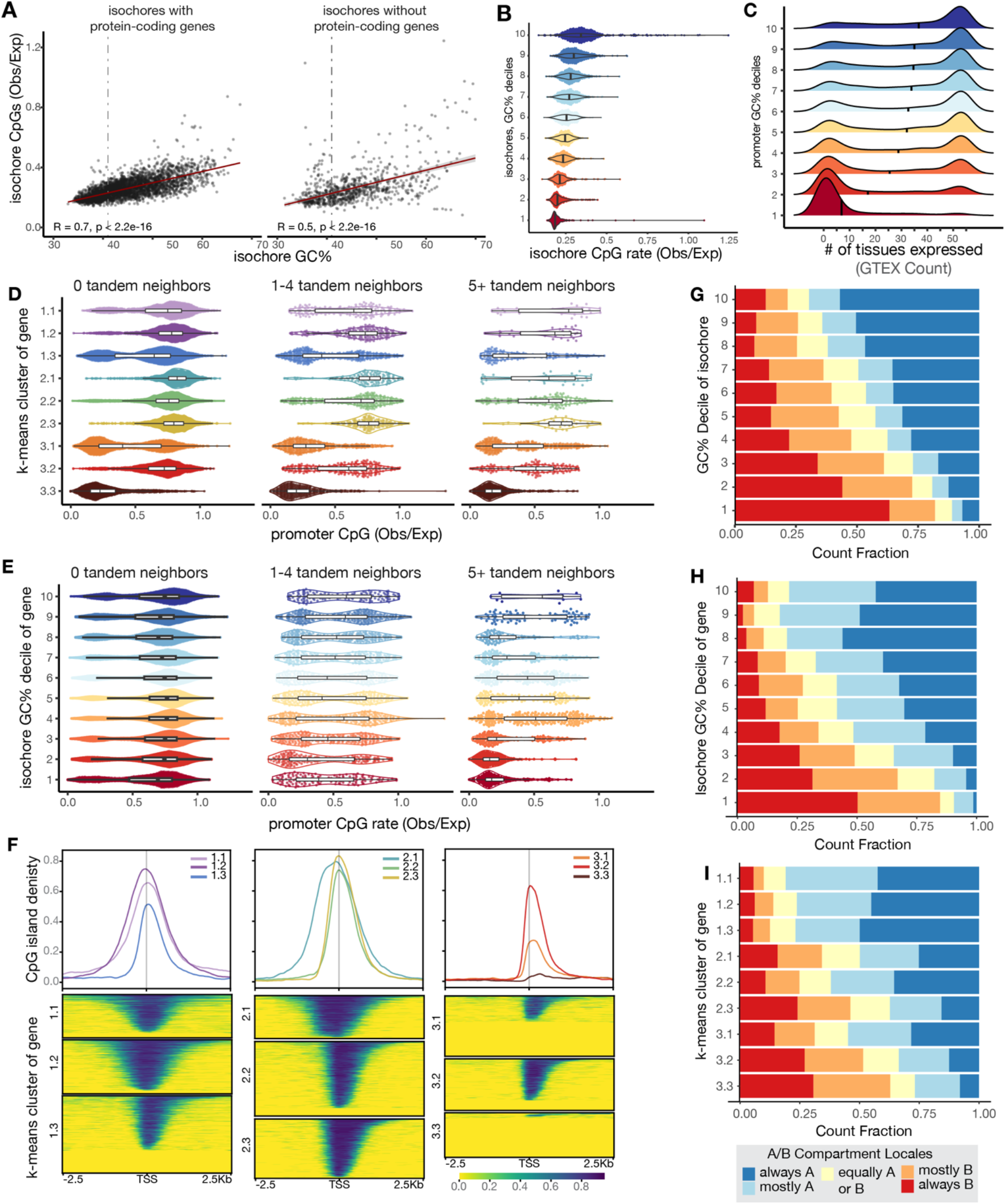
(A) Total CpG dinucleotide rate across isochores by their GC% content, split by isochores containing protein-coding genes and isochores without protein-coding genes. (B) Violin plots displaying total CpG rate across isochores, binned by decile. (C) Ridgeline plots displaying GTEX tissue expression distribution for genes binned by promoter GC% decile. (D) Distribution of promoter CpG rate of genes binned by k-means cluster assignment, split by degree of tandem duplication. (E) Distribution of promoter CpG rate of genes binned by isochore GC% decile, split by degree of tandem duplication. (F) CpG island density around the TSS (grey line) for genes in each k-means cluster. CpG island density reflects strength of called CpG islands from USCS unmasked CpG islands (54). Mean CpG rates across gene TSS in each cluster are summarized in line plots, while heatmaps represent island calls around the TSS for each gene. (G-I) Hi-C compartment assignment across 21 tissues (96) for isochores (G), or genes in each isochore (H) or k-means cluster (I). “Always A” and “always B” means the gene was assigned to that compartment in every sampled tissue.

